# Stability vs flexibility: reshaping archaeal membranes in silico

**DOI:** 10.1101/2024.10.18.619072

**Authors:** Miguel Amaral, Felix Frey, Xiuyun Jiang, Buzz Baum, Anđela Šarić

**Affiliations:** Institute of Science and Technology Austria, Klosterneuburg, Austria; Department of Physics and Astronomy, University College London, London, UK; MRC Laboratory of Molecular Biology, Cambridge, UK

## Abstract

Cellular membranes differ across the tree of life. In most bacteria and eukaryotes, single-headed lipids self-assemble into flexible bilayer membranes. By contrast, thermophilic archaea tend to possess bilayer lipids together with double-headed, monolayer spanning bolalipids, which are thought to enable cells to survive in harsh environments. Here, using a minimal computational model for bolalipid membranes, we explore the trade-offs at play when forming membranes. We find that flexible bolalipids form membranes that resemble bilayer membranes because they are able to assume a U-shaped conformation. Conversely, rigid bolalipids, which resemble the bolalipids with cyclic groups found in thermophilic archaea, take on a straight conformation and form membranes that are stiff and prone to pore formation when they undergo changes in shape. Strikingly, however, the inclusion of small amounts of bilayer lipids in a bolalipid membrane is enough to achieve fluid bolalipid membranes that are both stable and flexible – resolving this trade-off. Our study suggests a mechanism by which archaea can tune the material properties of their membranes as and when required to enable them to survive in harsh environments and to undergo essential membrane remodelling events like cell division.

---

A bounding membrane separates the interior of every biological cell from its extracellular environment and gives the cell its shape. To perform its function, this bounding membrane must be sturdy and impermeable to the leakage of small charged molecules like ions to allow the generation of an electrochemical gradient [1], but also flexible enough to be remodelled during essential cellular processes like cell division and vesicle formation [2]. These requirements are partially conflicting and difficult to combine.

Interestingly, two different generic membrane designs have evolved across the tree of life to solve this problem [3]. Bacterial and eukaryotic membranes possess fatty acid lipids that have a hydrophilic head group linked to hydrophobic tails via ester-linkages, which self-assemble into bilayer membranes [4]. By contrast, archaeal membranes lipids are constructed from branched isoprenoid lipids. These can include cyclopentane rings and are attached via an ether linkage to one (e.g., archaeol, Fig. 1A left) or two hydrophilic heads, leading them to be called bipolar lipids or simply bolalipids (e.g., caldarchaeol, Fig. 1A right) [4]. The hydrophilic heads can be composed of different functional groups with phosphatidyl and sugar being the most relevant moieties. For bolalipids the two head groups at either end of the molecule are typically distinct (Fig. 1A right) [5]. As an ensemble, bilayer and bolalipids can self-assemble into fluid lipid membranes of different architectures – as bilayer lipid membranes, bolalipid membranes or mixture membranes (Fig. 1B right).

**Figure 1.**
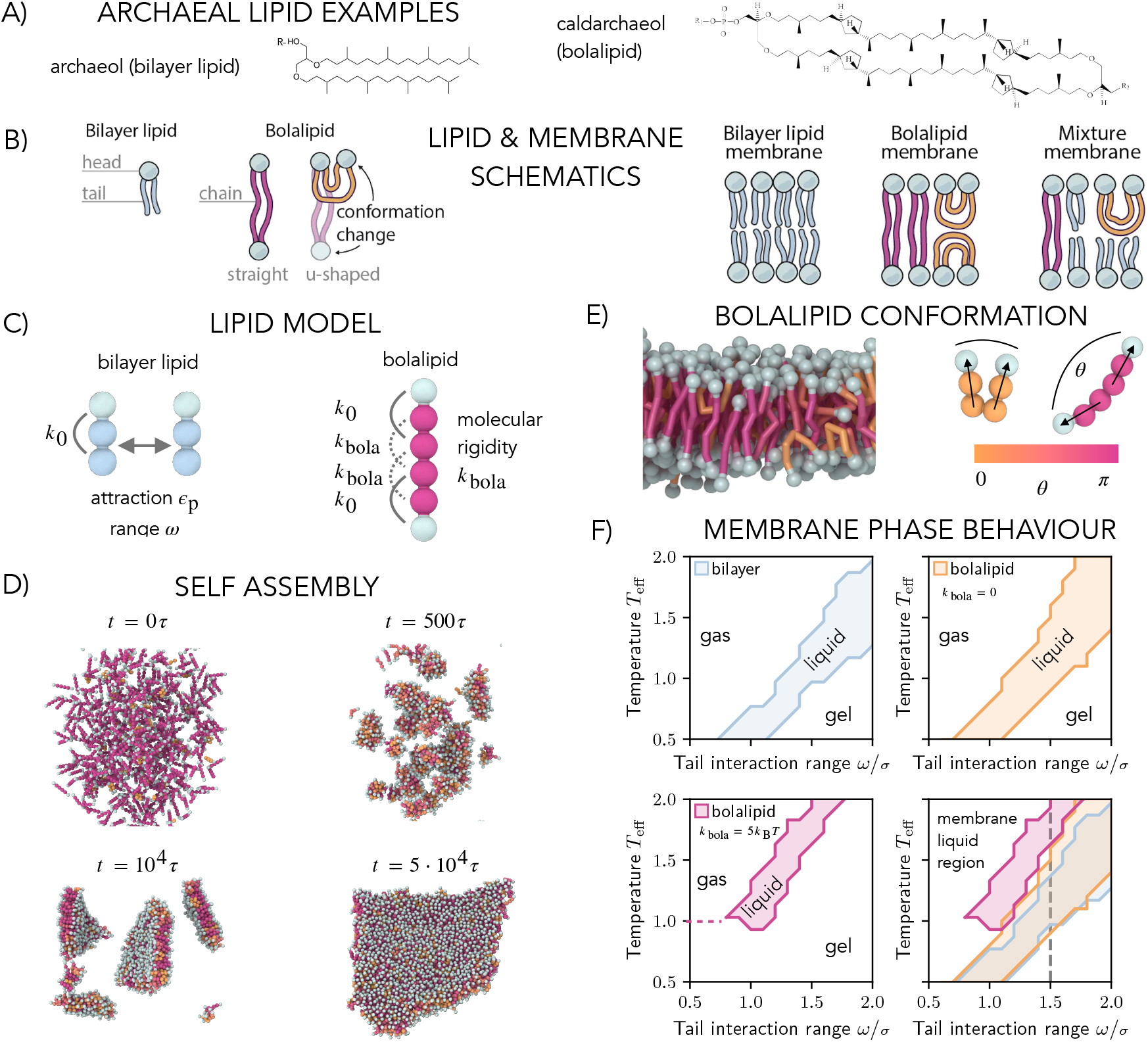
Computational model and phase space of bilayer and bolalipid membranes. (A) Structure of the diether bilayer lipid archaeol (left) and the tetraether bolalipid caldarchaeol including four cyclopentane rings (right), both present in the membrane of *Sulfolobus acidocaldarius*, a common archaeal model system that lives at high temperatures and low pH [6]. The hydrophilic head of a bolalipid can be composed of different functional groups represented by R1 and R2 (right). (B) Schematics for a bilayer lipid, a bolalipid and its two in-membrane conformations, membranes made of bilayer molecules only, bolalipid molecules only, and a mixture of the two (left to right). (C) Bilayer lipid (left) is described with one head bead and two tail beads straightened by an angular potential of strength *k*_0_. Tail beads of different lipids attract with the strength *ϵ*_p_ and the range *ω*. Bolalipids (right) consist of two bilayer lipids connected by a bond and straightened by an angular potential of strength *k*_bola_. (D) Snapshots of bolalipids self-assembling into a flat membrane (*k*_bola_ = 0.3 *k*_B_*T*). (E) Cross-section of self-assembled membrane (right), with bolalipids coloured according to their conformation: straight lipids in crimson and U-shaped lipids in orange. (F) Membrane phase behaviour: liquid, gel and gas regions as a function of the effective temperature *T*_eff_ and tail interaction range *ω* for bilayer (top left) and membranes made of flexible *k*_bola_ = 0 (top right) and stiff *k*_bola_ = 5 *k*_B_*T* (bottom left) bolalipid molecules. Overlays of all liquid regions (bottom right) show that stiffer lipids exhibit fluid membrane region at higher temperatures. The dashed line marks *ω* = 1.5*σ*, the value we used in the rest of the work.

Like for bacteria and eukaryotes, archaea must keep their lipid membranes in a fluid state (homeoviscous adaptation). This is important even under extreme environmental conditions, such as hot and cold temperatures, or high and low pH values [7]. Because of this, many archaea adapt to changes in their environment by tuning the lipid composition of their membranes: altering the ratio between bola- and bilayer lipids in their membranes [8, 9] and/or by changing the number of cyclopentane rings in their lipid tails, which are believed to make lipid molecules more rigid [5]. For example, *Thermococcus kodakarensis* increases its tetraether bolalipid ratio from around 50% to over 80% when the temperature of the environment increases from 60 to 85°C [10]. Along the same lines, the cell membrane of *Sulfolobus acidocaldarius*, can contain over 90 % of bolalipids with up to 8 cyclopentane rings at 70°C and pH 2.5 [5, 11]. It is worth mentioning that in exceptional cases bacteria also synthesise bolalipids in response to high temperatures [12], highlighting that the study of bolalipid membranes is relevant not only for archaeal biology but also from a general membrane biophysics perspective. Besides temperature adaptation, archaeal lipids also exhibit dense packing, high viscosity, and low porosity to small molecules like protons [10].

All this makes the study of reshaping archaeal membranes extremely interesting. Unfortunately, it is fairly challenging to investigate archaeal cells experimentally due to their unusual chemistry and extreme living conditions [13]. Therefore, most of what we know about archaeal tetraether lipid membranes thus far has been collected from studying *in vitro* reconstituted membranes. For instance, the conformation of individual lipids in bolalipid membranes was studied at the water-air interface [14, 15] or using NMR experiments on lipid vesicles [16]. This suggested the existence of U-shaped lipid conformations in archaeal type membranes (Fig. 1B). Moreover, *in vitro* reconstituted vesicles primarily composed of bolalipids can fuse with influenza virus particles at similar kinetic rates compared to bilayer vesicles, further suggesting that bolalipids exist in U-shape allowing for membrane remodelling and fusion [17]. Membrane properties like bending stiffness [18] or lipid phase [19] have also been measured in vesicles prepared from archaeal tetraether lipids to demonstrate that archaeal lipids derived membranes are exceptional in being stable up to temperatures of 80°C [19]. Moreover, experiments with lipid vesicles made from synthetic bilayer lipids that include cyclopentane rings, which naturally appear in lipids of extremophilic archaea, showed that increasing the number of these rings increases membrane rigidity [20].

From the view point of membrane physics, the remodelling of bilayer membranes has been studied for decades using continuum models and computer simulations [21–23]. By contrast, only a few studies have investigated bolalipid membranes applying computational or theoretical tools [24, 25]. Specifically, the pore closure time in bolalipid membranes, and the role of cyclopentane rings for membrane properties has been investigated using all-atom simulations, showing decreased lateral mobility, reduced permeability to water, and increased lipid packing [26–28]. Moreover, using coarse-grained simulations, it was suggested that bolalipid membranes are thicker [29], exhibit a gel-to-liquid phase transition at higher temperature [30], and exhibit a reduced diffusivity [31]. However, little research has been devoted to investigating mechanics and reshaping of bolalipid membranes at the mesoscale despite the obvious importance of this question from evolutionary, biophysics, and biotechnological perspectives and although different membrane physics is expected to manifest.

Here we have designed a minimal model of archaeal membranes that can be used to explore the impact of both bolalipids and bilayer lipids on membrane biophysics and membrane reshaping. Our coarse-grained molecular dynamics simulations show that the geometry of bolalipids is, through the effects of entropy alone, sufficient to shift the fluid phase of archaeal-type membranes so that they are stable at high temperatures. We show that membranes assembled from bolalipids can have a much higher bending rigidity than bilayer-derived membranes, and are more likely to resist membrane shape changes as in fission. During membrane deformation, stress in these bolalipid membranes is relieved by a small fraction of bolalipids taking up a U-shaped conformation, which renders them a mechanically switchable material. Without these U-shaped bolalipids, stress leads to the formation of large pores. However, when combined, a mixture of bilayer and bolalipids generates membranes that are stable at high temperature, which become softer as the fraction of bilayer lipids increases. Taken together, our results demonstrate how doping a bolalipid membrane with a small fraction of bilayer lipids can relieve the trade-off, enabling membranes to be both stable under extreme conditions and to be remodelled without leaking.

## RESULTS

### Computational Model

To study bolalipid membranes and compare them to bilayers, we extended the Cooke and Deserno model for bilayer membranes [32]. In the original model for the bilayer a single bilayer lipid is represented by a chain of three nearly equally sized beads of diameter of ~ 1*σ*, where *σ* is our distance unit and roughly maps to 1 nm (Fig. 1C left); one bead stands for the head group (cyan) while the others represent the hydrophobic tail (blue). Each adjacent pair of beads in a lipid is linked by a finite extensible nonlinear elastic (FENE) bond. The angle formed by the chain of three beads is kept near 180° via an angular potential with strength *k*_0_, instead of the approximation by a bond between end beads of the original model [32].

While lipid heads interact exclusively through volume exclusion, the beads of lipid tails interact via a soft attractive potential of the strength *c*_p_ and range *ω* (Fig. 1C left), effectively modelling hydrophobic interaction in an implicit solvent. This interaction strength governs the membrane phase behaviour and can be interpreted as the effective temperature or reduced temperature *T*_eff_ = *k*_B_*T* /*ϵ*_p_. As the distinction between scaling interactions (*T*_eff_) or temperature (*T*) is not important for our analysis (see Supplemental Information (SI) section 14), for simplicity we refer to *T*_eff_ as temperature in the following.

To model a bolalipid molecule, we joined two bilayer lipids so that a lipid molecule is formed with a head bead (cyan) that is linked to four tail beads (crimson) which are again linked to another head bead (Fig. 1C right). In this way, both bilayer lipids and bolalipids share the same molecular structure and the same interactions between lipid beads. To decouple the effect of the connected geometry of the bolalipids from that of lipid asymmetry, we assume both head beads of a bolalipid to share the same properties. Bolalipids in archaeal membranes can differ in the number of cyclopentane rings or the branching of the tail and thus in the molecular stiffness [7, 33]. To represent this effect, we added two angular potentials between the second and the fourth and the third and the fifth tail bead with variable strength *k*_bola_. By varying *k*_bola_, we can control the molecular stiffness of the bolalipid molecules and thus model different types of bolalipids. The model is simulated within molecular dynamics implemented in the LAMMPS open source package [34], and simulations are visualized with OVITO [35]. To include the implicit effect of the surrounding water and to simulate membranes at vanishing tension, we used a Langevin thermostat combined with a barostat that kept the membrane at zero pressure in the *x*-*y* plane. Details on the computational model are given in SI section 1.

### Self-assembly and phase behavior

We first tested the ability of both the bilayer and bolalipid molecules to self-assemble into membranes. To do so, we placed dispersed lipids in a periodic 3D box. For both bilayer and bolalipids we found that membranes self-assembled over a wide range of lipid interaction parameters (Fig. 1D, SI section 2, and Movie S1). We then explored the influence of flexibility in the bolalipid molecule. At small values of the molecular rigidity *k*_bola_, single lipids are flexible and can thus adopt a range of possible conformations. The different conformations can be classified by the angle *θ* between the two lipid heads (Fig. 1E). Two lipid conformations dominated the conformation distribution in the context of a membrane: the U-shaped conformation with both head beads on the same membrane leaflet (*θ* ≈ 0) and the straight conformation with one head bead in each opposing membrane leaflet (*θ* ≈ *π*) (Movie S2.b, Fig. S1, and SI section 3; for comparison, bilayer membranes in Movie S2.a and rigid bolalipid membranes in Movie S2.c). In the self-assembled bolalipid membrane, we marked bolalipids as being in the U-shape conformation if *θ < π*/2 and in the straight conformation otherwise.

To study the phase behaviour of bilayer and bolalipid membranes, we analysed the diffusion of single lipids as a function of the lipid interaction parameters *ϵ*_p_ and *ω* (SI sections 4 and 5). The diffusion constant *D* exhibited a discontinuity as a function of the temperature, which marks the transition as the membrane moves from the gel phase to the liquid phase. The discontinuity occurred at different values of interaction strength *ϵ*_p_ and interaction range *ω* for bilayer and bolalipid lipids, and it also depended on the values of the molecular stiffness *k*_bola_ (Fig. S4). In these simulations, the disintegration of the membrane defined an upper limit to the liquid phase and the transition to the gas phase. Based on this classification, we plotted the phase diagram for bilayer membranes (Fig. 1F top left), fully flexible bolalipid membranes (*k*_bola_ = 0, Fig. 1F top right), and rigid bolalipid membranes (*k*_bola_ = 5 *k*_B_*T*, Fig. 1F bottom left) as a function of the range of the hydrophobic interaction *ω* and the temperature *T*_eff_. Membranes made of bilayers (blue) and flexible bolalipid molecules (*k*_bola_ = 0, orange) behaved similarly under these conditions (Fig. 1F bottom right). Strikingly, just as observed in extremophile archaea, as the molecular stiffness of bolalipids increased (*k*_bola_ = 5 *k*_B_*T*, magenta), the liquid region was shifted toward higher temperatures and larger values of the interaction range. This is due to the fact that bolalipid molecules are able to engage in more extensive interactions with partners when in the extended conformation, which helps to stabilize the membrane at higher temperatures.

### Bolalipid conformations and mechanical properties

To explore mechanical properties of bolalipid membranes, we chose *ω* = 1.5 *σ* for the remainder of the work (dashed line in Fig. 1F bottom right). Fig. 2A shows the phase diagram for bolalipid membranes replotted as a function of temperature and bolalipid rigidity for the chosen interaction range. To be able to compare membranes made of bolalipids of different molecular rigidities, we needed to adjust temperature for each to reach similar fluidities, shown by dashed lines in Fig. 2A. We then characterised the conformations of individual bolalipids in flat membranes, measured by the fraction of bolalipids in U-shape conformation *u*_f_. We find that for flexible bolalipids (*k*_bola_ = 0), more than 50% of all lipids are in the U-shape conformation (Fig. 2B). This fraction decreases with increasing rigidity of bolalipid molecules and vanishes around *k*_bola_ *≥* 2 *k*_B_*T*, for which almost all bolalipids take up linear conformations.

**Figure 2.**
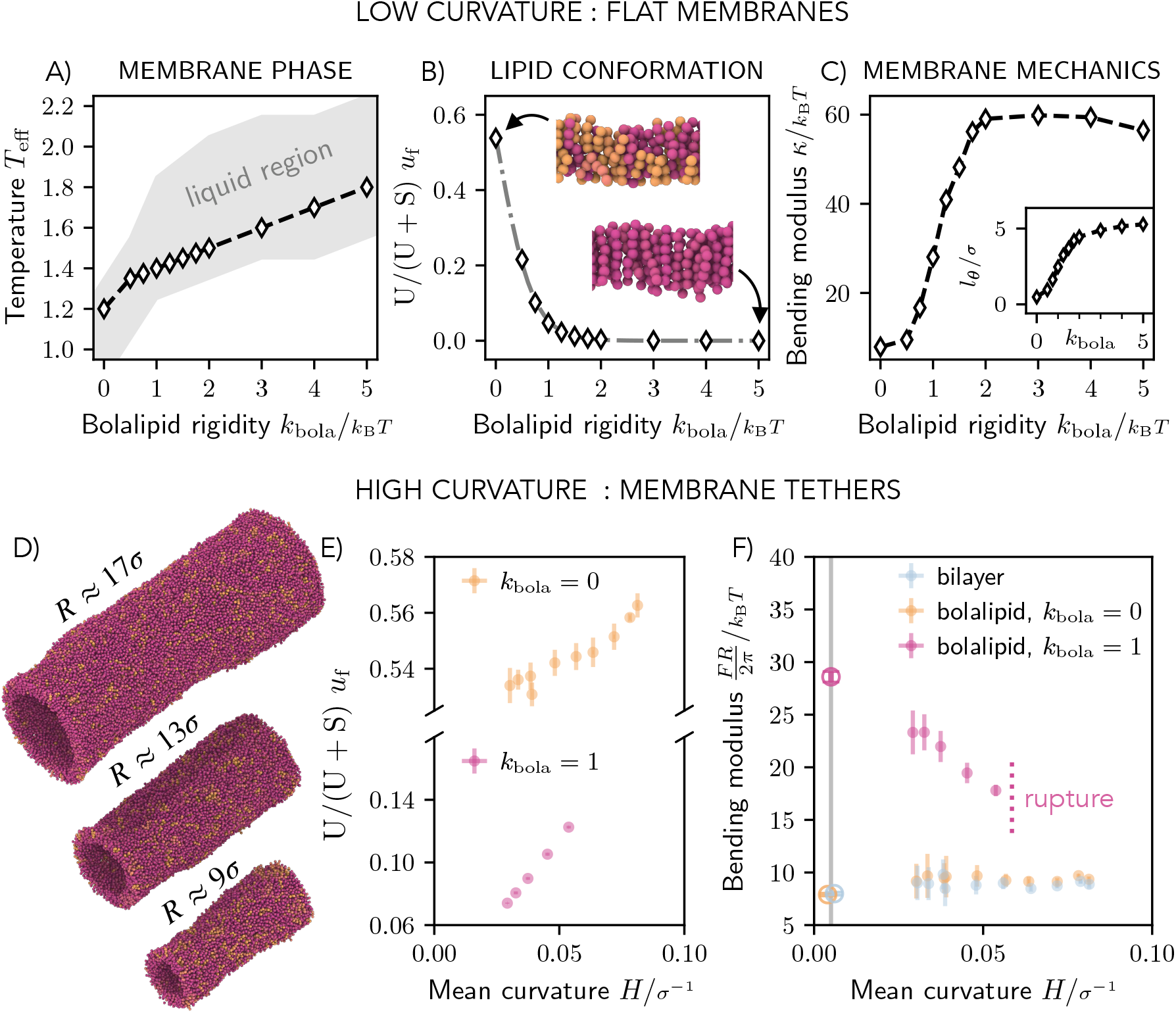
Mechanics of pure bolalipid membranes. (A) Liquid region as a function of temperature and bolalipid rigidity for pure bolalipid membranes (gray). The dashed line shows the bolalipid membranes of approximately same fluidities. (B) Fraction of bolalipids in the U-shape conformation (*θ* = 0), fitted to *u*_f_(*k*_bola_) = 1/(1 + exp(*β*(−0.16 + 3*k*_bola_))) (gray dashed line) according to a two-state model. Insets: simulation snapshots with bolalipids coloured according to their conformations. (C) Bending modulus as a function of bolalipid molecule rigidity *k*_bola_. Inset: Tilt persistence length *l*_*θ*_ as a function of bolalipid rigidity *k*_bola_. (D) Snapshots of bolalipid membranes at the range of explored curvatures for *k*_bola_ = 1*k*_B_*T*. (E) Fraction of bolalipid molecules in the U-shaped conformation as a function of the mean membrane curvature *H* = 1/(2*R*) for membranes made of flexible (*k*_bola_ = 0) and semi-flexible (*k*_bola_ = 1*k*_B_*T*) bolalipid molecules. (F) Bending modulus as a function of curvature. For the flat membrane (*H* ≈ 0), the corresponding bending rigidity from (C) is marked by the vertical line and empty circles.

This behaviour can be easily captured by considering bolalipids as a two state system, with straight and U-shaped conformations, as argued before (Fig. S1). We assumed that the energy of the straight conformation vanishes *E*_s_ = 0 and the energy of the U-shaped conformation reads *E*_u_ = *c*_0_ + *c*_1_*k*_bola_, where, for simplicity, we assumed that the energy linearly depends on *k*_bola_ which is the only relevant energy scale in the system and the two constants *c*_0_ and *c*_1_, which determine the conformation energy of U-shaped bolalipids. The fraction of bolalipids in U-shape conformation then follows *u*_f_(*k*_bola_) = 1/(1 + exp(*β*(*c*_0_ + *c*_1_*k*_bola_))), with *β* = 1/(*k*_B_*T*), as shown by the fit in Fig. 2B (dashed gray line, *R*^2^ = 0.99, see SI section 6). For the fit it appears that *c*_0_ *<* 0, which implies that bolalipids in U-shape conformation are slightly favoured over straight bolalipids at *k*_bola_ = 0 (explored in SI section 6).

### Bolalipid conformations and membrane rigidity

It was previously hypothesized that an increasing fraction of bolalipids in straight configuration would increase the membrane rigidity [18]. To determine the membrane rigidity using our model, we assessed the height fluctuation spectrum (*h*^2^) of flat membranes in a periodic box. [32, 36]. Interestingly, we found that the original theory of Helfrich [37] failed to describe the resulting height fluctuation spectrum. However, the extended theory by Hamm and Kozlov [38], which also includes the energetic cost of lipid tilt, successfully captured bolalipid fluctuations (SI section 16). In this case, the resulting height spectrum of the membrane at vanishing membrane tension is given by [36]

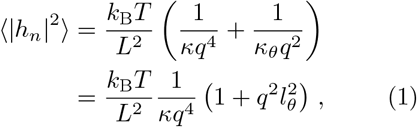

where *q* = 2*πn*/*L* is the wave number (*n ∈* ℤ), *L* is the box size, *κ* is the bending rigidity of the membrane, *κ*_*θ*_ is the tilt modulus and 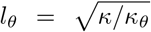 is a characteristic length scale related to tilt, which we call the tilt persistence length. Considering Eq. (1), the tilt term is expected to matter if the analysed inverse wave numbers become of the same order as the tilt persistence length *l*_*θ*_. The wave numbers that we analysed correspond to wavelengths that are at least twice the thickness of the membrane (*q <* 2*π*/(12*σ*) ≈ 0.5 *σ*^−1^). Thus, the tilt term is expected to noticeably contribute if *l*_*θ*_ *>* 2 *σ*. For typical bilayer membranes, one finds *κ*_*θ*_ = 12 *k*_B_*T* nm^−2^ [36] and *κ* = 20 *k*_B_*T* [39], so *l*_*θ*_ ~ 1 nm ≈ 1 *σ* and therefore the tilt term can be neglected as practised before [32]. However, when the membrane rigidity increases as we expect for bolalipid membranes, the tilt persistence length *l*_*θ*_ increases and the tilt term in Eq. (1) becomes relevant.

By fitting the height spectrum for bolalipid membranes (and bilayer membranes for comparison) (Fig. S6), we measured the bending rigidity (Fig. 2C) and the tilt persistence length *l*_*θ*_ (Fig. 2C inset) as a function of *k*_bola_ (see SI section 7 and Movie S3). With increasing bolalipid molecular rigidity *k*_bola_, the bending rigidity *κ* rose from 8 *k*_B_*T* and plateaued at 60 *k*_B_*T*, showing bolalipid membranes can be very rigid while liquid. Strikingly, the increase in membrane rigidity coincided with U-shaped bolalipids vanishing from the membrane (Fig. 2B), which confirmed the hypothesis that straight bolalipids render lipid membranes rigid. At the same time, the tilt persistence length *l*_*θ*_ increases with bolalipid rigidity, starting near zero, crossing the 2 *σ* threshold at *k*_bola_ = 1 *k*_B_*T* and plateauing at 5 *σ*. Since membrane bending and lipid tilting are two modes of membrane deformations that compete, we conclude that bilayer and flexible bolalipids molecules form flexible membranes that prefer to bend rather than to tilt, while bolalipids in straight configuration form rigid membranes that prefer to tilt rather than to bend. Taken together, bolalipid membranes made of flexible lipid molecules are as flexible as lipid bilayers, adopting U-shaped conformations, where those made of bolalipids in straight configurations are rigid.

### Interplay between membrane curvature and rigidity

Changing membrane curvature alters the area differently in the two membrane leaflets. To adapt to the area difference, we thus expect the fraction of U-shaped bolalipids to change as the membrane curvature changes. Moreover, the results of Fig. 2B and Fig. 2C showed that the U-shaped bolalipid fraction and the membrane bending rigidity are correlated. As a result, we predict that the fraction of straight versus U-shaped bolalipids in a membrane will change in response to membrane bending, in a way that makes the bending rigidity of a bolalipid membrane curvature dependent.

To investigate the effect, we measured the bending rigidity of bolalipid membranes in cylindrical shapes as a function of their radii [40] (see Fig. 2D, Movie S4, and SI section 8). We first noticed that while membrane tubes made of bilayer and flexible bolalipids were stable up to small cylinder radii *R*, almost as small as the membrane thickness itself, we found that membrane made from stiffer bolalipids (*k*_bola_ = 1*k*_B_*T*) ruptured well before. Strikingly, while for stiffer bolalipids (*k*_bola_ = 1*k*_B_*T*) the U-shaped bolalipid fraction increased strongly over a short range of the mean membrane curvature (*H* = 1/(2*R*)), we only found a small change in the U-shaped bolalipid fraction of flexible bolalipid membranes (*k*_bola_ = 0) (Fig. 2E). Consequently, since the fraction of U-shaped bolalipid molecules controls the bending rigidity of bolalipid membranes (Figs. 2B and 2C), we found that there is a strong dependency of the bending rigidity *κ* on the membrane mean curvature of stiffer bolalipids (*k*_bola_ = 1*k*_B_*T*) (Fig. 2F).

In an elastic material, the strain modulus holds constant and deformation is reversible. For bolalipid membranes at *k*_bola_ = 1*k*_B_*T*, however, the bending modulus decreases when deformation increases, rendering bolalipid membranes hypoelastic. Fortunately, this dependency on curvature does not invalidate our fluctuation results, where the curvature is small enough that its effect on the bending modulus is negligible (SI section 15). In contrast, we did not find that *κ* was curvature-dependent for bilayer or flexible bolalipid membranes (*k*_bola_ = 0). Taken together, the rigidity of bolalipid membranes is not only controlled by the molecular stiffness of their lipid constituents but also by the emerging geometry of the ensemble of lipids. Since membrane geometry and thus membrane rigidity will change upon membrane deformations this gives rise to hypoelastic material properties.

### Gaussian rigidity of bolalipid membranes

Another important material parameter is the Gaussian bending modulus 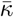, which characterizes the reshaping behaviour of fluid lipid membranes under topological changes [39]. 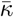 is notoriously difficult to measure since it only becomes detectable when the membrane changes its topological state. Continuum membrane theory, combining shape stability arguments and elasticity, predicts 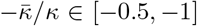 [39], where the former value is expected for incompressible membranes. Indeed, most of the numbers we know for the ratio of the two bending rigidities, many of which were deduced from simulations, lie within this range [41]. Using the same method as developed by Hu et al. [41], we determined 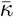 by measuring the closing efficiency of membrane patches into a sphere (see SI section 9). At *T*_eff_ = 1.2, we obtained 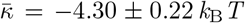 and thus a ratio of 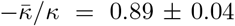 for bilayer membranes, similar to what has been reported previously [41]. For flexible bolalipid membranes, we got a slightly smaller value for 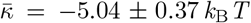. Due to the larger bending modulus, however, flexible bolalipid membranes show a significantly smaller ratio 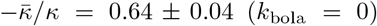. At larger temperature (*T*_eff_ = 1.3), the ratio can be even smaller 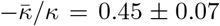 (see SI section 9). The result shows that in addition to the differences in bending rigidities, the ratio of the two bending moduli differs strongly in bilayer and bolalipid membranes.

### Archaeal membranes made of mixtures of bolalipids and bilayer-forming lipids

Archaeal membranes contain varying amounts of bilayer lipids [5, 8]. The exact bolalipid/bilayer fraction depends on the growth temperature, with higher levels of bolalipids with increasing temperature [9], and higher fraction of cyclopentane rings in the tails [33]. In order to investigate the effect of different lipid contents on membrane mechanical properties, we wanted to model the archaeal membrane by mixing bilayer lipids into bolalipid membranes. Since in our model, the liquid regions of rigid bolalipid membranes and bilayer membranes do not overlap (Fig. 1F bottom right), we picked the temperature *T*_eff_ = 1.3 to minimize fluidity mismatch and we set the molecular rigidity *k*_bola_ = 2 *k*_B_*T* to limit U-shaped bolalipids (Fig. 2B). We then measured the diffusion constant *D* as a function of the fraction of bilayer lipids *f* ^bi^ (see SI section 5). Interestingly, we found that mixing only 10% bilayer lipids into the bolalipid membrane in gel state is enough to fluidize the membrane (Fig. 3A). We then measured the bending rigidity and the tilt persistence length *l*_*θ*_ of flat mixture membranes by analysing the fluctuation spectrum. Both the bending rigidity *κ* (Fig. 3B) and the tilt persistence length *l*_*θ*_ decreased non-linearly with the bilayer lipid fraction *f* ^bi^ (Fig. 3B inset). Taken together, the bolalipid membrane can be substantially softened either through bolalipids acquiring the U-shaped conformation or through addition of bilayer-forming lipids.

**Figure 3.**
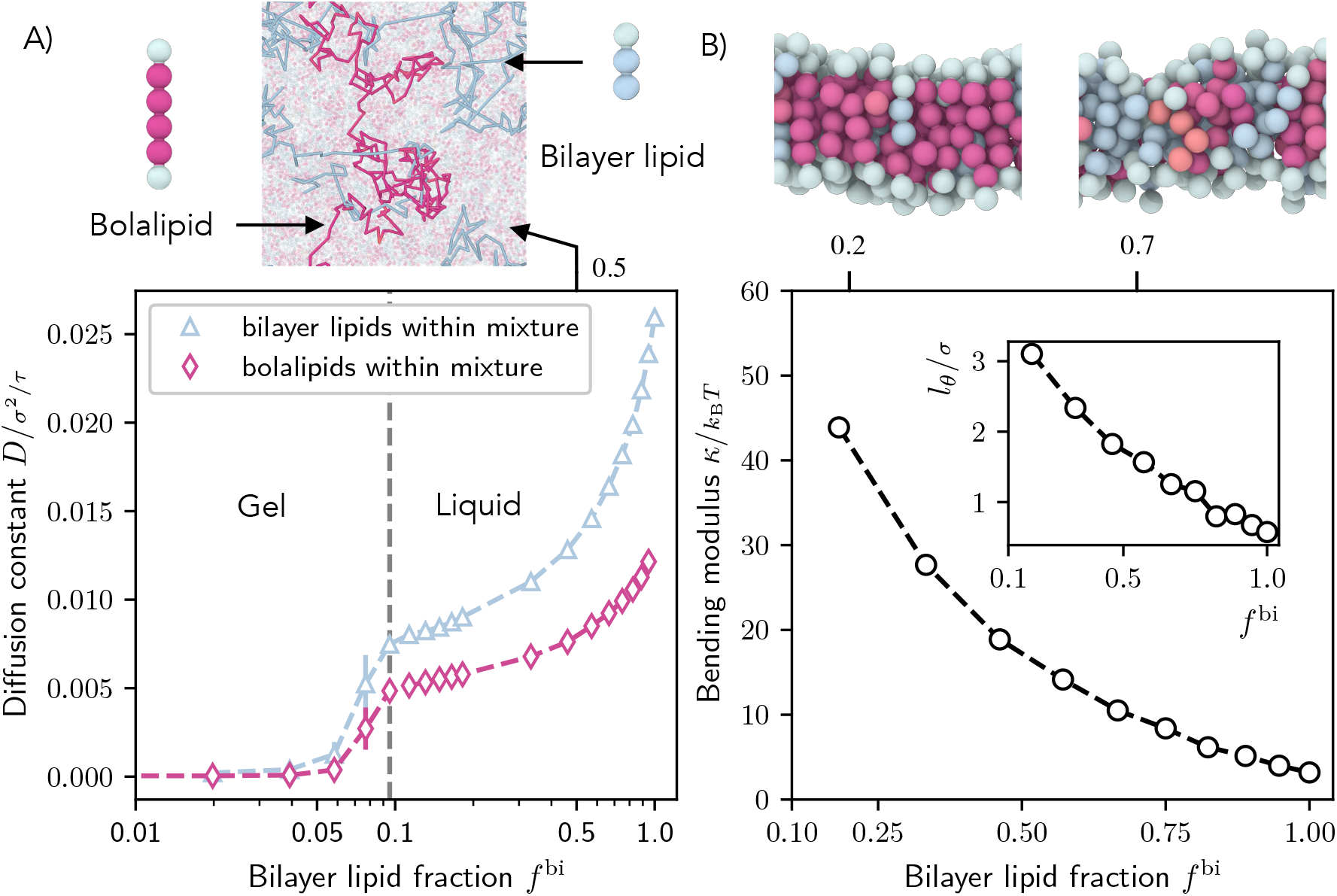
Fluidity and rigidity of mixed bilayer/bolalipid membranes. (A) Single lipid diffusion constant for each species as a function of bilayer fraction *f* ^bi^ (*k*_bola_ = 2*k*_B_*T*, *T*_eff_ = 1.3). For *f* ^bi^ *≥* 0.1, the resulting mixture becomes liquid. Top: Diffusion trajectories of a bolalipid (crimson) and a bilayer lipid (blue) in a mixture membrane at *f* ^bi^ = 0.5. (B) Bending rigidity *κ* and (Inset) tilt persistence length *l*_*θ*_ as a function of the fraction of bilayer molecules *f* ^bi^. Top: Snapshots show bilayer lipids (blue) in mixed membranes at two different values of *f* ^bi^.

### Curving bolalipid membranes

To investigate the response of bolalipid membranes to large membrane curvature and topology changes like those induced upon vesicle budding, which regularly occurs in archaea, we simulated membrane wrapping of an adhesive cargo bead (Fig. 4A, Movies S5.a to S5.c, and SI section 10). Importantly, this provided us with a method to study how lipid organization is affected by externally imposed membrane curvature and mechanics. We first simulated membrane wrapping at different adhesion energies *ϵ*_mc_ between lipid head beads and the cargo until we observed that the membrane wrapped the cargo completely (including membrane fission). Then the minimum adhesion energy, for which a membrane bud completely enveloped the cargo bead, is the onset adhesion energy 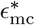 (Fig. 4A), which we measure as a function of the bolalipid stiffness *k*_bola_ (Fig. 4B). For small molecular stiffness 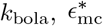 first increases linearly with *k*_bola_ before it saturates around *k*_bola_ = 3 *k*_B_*T*. We expect that the onset energy is proportional to the membrane bending rigidity 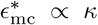, because the bending energy to wrap a spherical particle is size-invariant [39, 42]. When we increased the bending rigidity, through increasing stiffness of bolalipid molecule 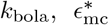 increased by a factor of 3 (Fig. 4B), suggesting that also *κ* increased by a factor of 3. However, from directly measuring the membrane rigidity from the fluctuation spectrum (Fig. 2C), we saw that *κ* increased by a factor of 10. To reconcile these seemingly conflicting observations we reason that the bending rigidity *κ*, similar to Fig. 2F, is not constant but softens in the range *k*_bola_ *∈* [0, 2]*k*_B_*T*, upon increasing membrane curvature. This is due to the dynamic change in the ratio between bolalipids in straight and U-shaped conformation. Hence, bolalipid membranes show marked hypoelasticbehaviour as they soften during reshaping.

**Figure 4.**
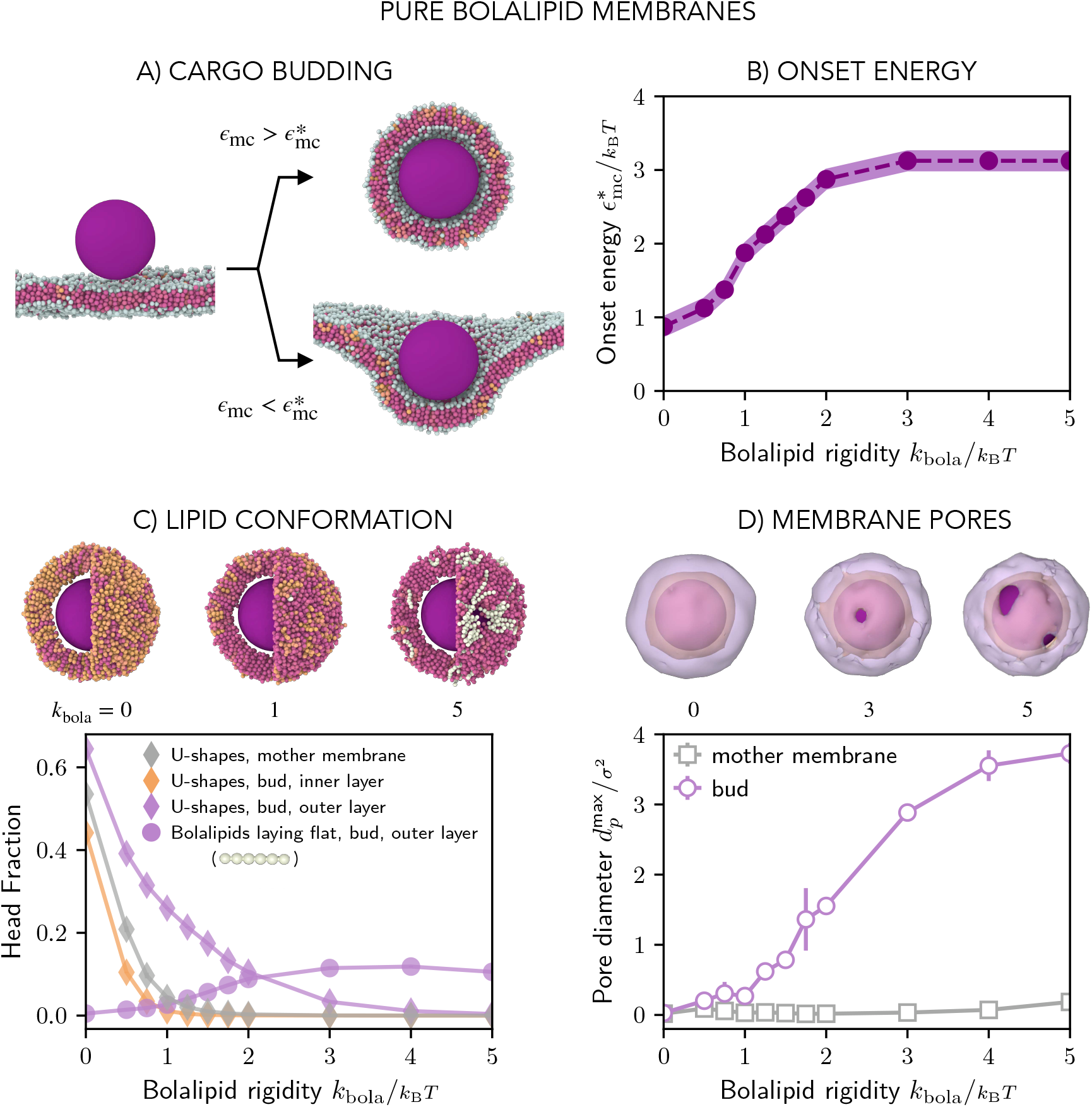
Reshaping of pure bolalipid membranes. (A) Simulation snapshots of the membrane wrapping a cargo bead adhering to it. Above the onset adhesion energy 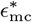, the cargo is fully wrapped by the membrane and buds off the mother membrane. (B) Onset energy 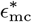 as function of the bolalipid molecule rigidity *k*_bola_ (for the parameters defined by the line in Fig. 2A). (C) Bottom: Fraction of bolalipids in the U-shape conformation *u*_f_ in the outer and inner layers of the membrane bud, and in the flat mother membrane, as function of the bolalipid molecule rigidity *k*_bola_. Top: Snapshots and cross-sections of the membrane around the cargo bud. At high bolalipid rigidity the pores form around the cargo, and are lined with bolalipid molecules lying flat around the pore in a straight conformation, with both heads in the outer layer (coloured in white). The rest of bolalipids coloured according to their head-to-head angle as before. (D) Bottom: Average diameter of transient pores in the membrane bud and the mother membrane as function of the bolalipid molecule rigidity *k*_bola_. Pores are defined as membrane openings through which a sphere of diameter 1*σ* can cross. Top: Snapshots of the membrane surface with outer and inner leaflet surface coloured in purple and orange, respectively, intersecting at the rim of the pore (gray).

Through analysing the bolalipid conformations, we found that the membrane was able to curve by increasing the fraction of U-shaped bolalipids in the outer layer of the deformation (Fig. 4C and SI section 12). To a lesser degree, we also observed this effect on the inner membrane neck (Fig. S14). Remarkably, though, even when lipids are so stiff that there are no more U-shaped bolalipids in the flat mother membrane (*k*_bola_ = 2 *k*_B_*T*), the outer layer of the a curved membrane retained a non-negligible fraction of ≈ 10% U-shaped bolalipids, which in turn decreases *κ* and softens the membrane. However, the softening effect on the membrane, indicated through a constant onset energy for *k*_bola_ *≥* 3 *k*_B_*T* (Fig. 4B), persists even for those very stiff bolalipids. Since for stiff membranes, practically all U-shaped bolalipids are gone (Fig. 4C), this suggested that an additional membrane-curving mechanism must be involved.

Looking more closely, at high molecular rigidity (*k*_bola_ *≥* 2*k*_B_*T*) we observed the formation of multiple pores on the membrane bud, which we quantified by measuring the time-averaged maximum pore diameter (Fig. 4D, see SI section 12 and Movie S6). While large pores were not observed in the flat membrane, the diameter of membrane pores around the cargo was found to grow with the increase in bolalipid stiffness. We reasoned that pores form when the energetic cost required to change the bolalipid conformation to release bending stress is larger than the energetic cost of opening a lipid edge surrounding the pore. Hence, for relatively flexible bolalipids, U-shaped bolalipids provide the necessary area difference between the outer and inner layer of the membrane bud and thereby soften the membrane. For stiff bolalipid molecules, however, membrane pores start to form to enable membrane curvature as U-shaped bolalipids become prohibited. Both mechanisms help to explain the discrepancy between 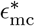 and the bending modulus *κ* obtained by studying membrane fluctuations (Fig. 2C).

### Curving archaeal membranes

Having shown that bolalipid membranes can effectively soften also by including some amount of bilayer-forming lipid molecules, we next measured the onset energy 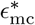 for cargo budding in the membranes formed by mixtures of bolalipids and bilayer-forming lipids, as a function of bilayer lipid head fraction 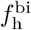 (Fig. 5A and Movie S5.d). We found that the onset energy sharply decreases with increasing amount of bilayer forming lipids, and plateaus for 50% bilayer head fraction, where it acquires similar values as in the case of fully-flexible bolalipids (Fig. 4B). For small bilayer fractions, U-shaped bolalipids localize almost exclusively on the outer layer of the bud (Fig. 5B). As the bilayer fraction increases, there is a steady reduction in the percentage of U-shaped bolalipids in the outer layer in favour of bilayer lipids that take their role in supporting membrane curvature, with U-shaped bolalipids completely vanishing at high fractions of bilayer-forming lipids. The fraction of bilayer lipid head beads initially shows an asymmetry between the preferred outer layer and the penalized inner layer around the cargo (Fig. 5C), but eventually approaches 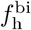 in both layers. Taken together, as the bilayer lipid fraction increases, the role of U-shaped bolalipids in making up the asymmetry between the outer and inner layer, is taken over by bilayer lipids. Curiously, the addition of bilayer lipids promotes the formation of U-shaped lipids, both in the flat membrane and even in the inner layer around the bud. This is likely to be explained by the fact that when more bilayer lipids are incorporated, the membrane is less densely packed (Fig. S12B) and thus U-shaped bolalipids are promoted.

**Figure 5.**
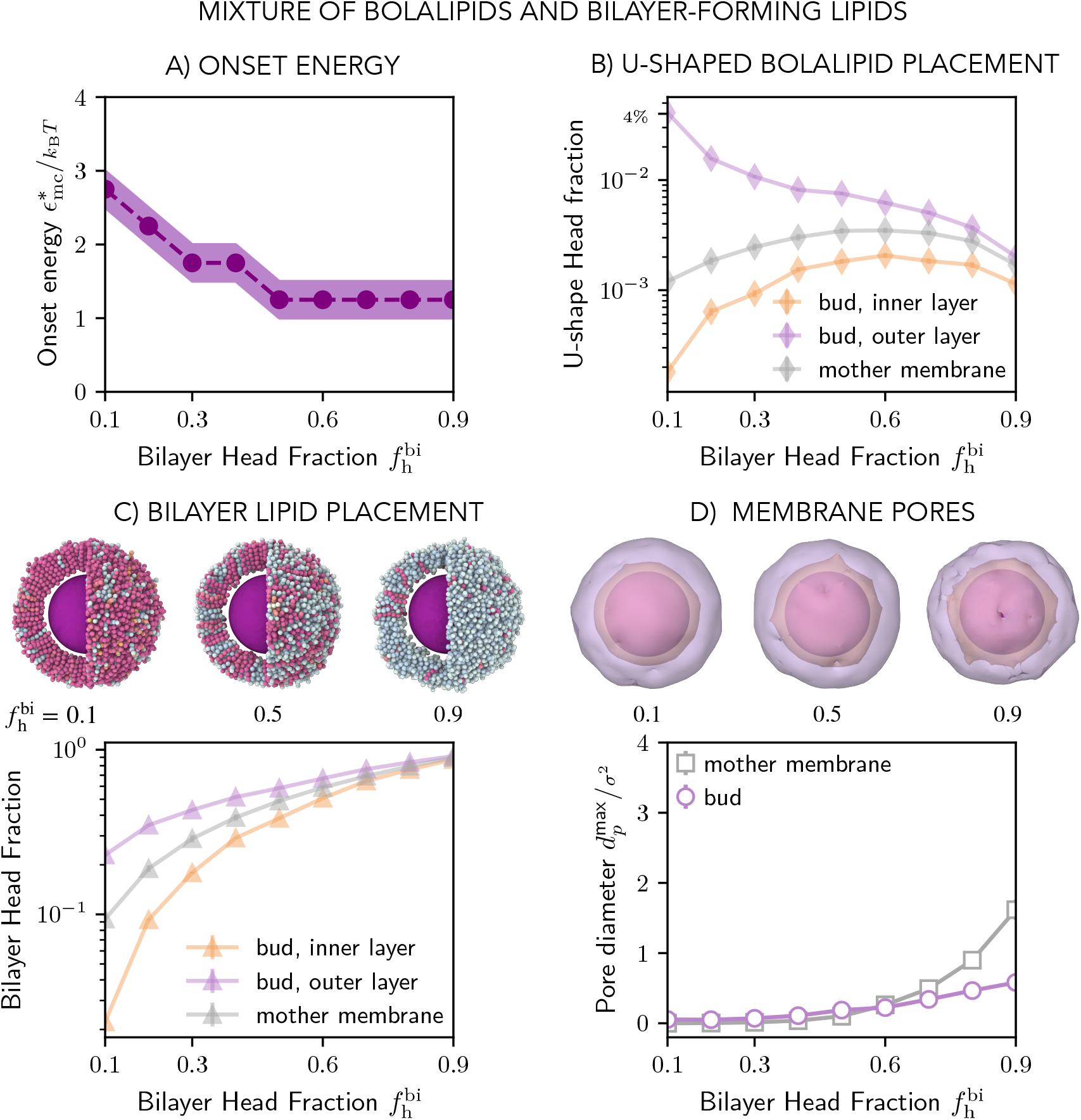
Curving of the mixed membranes, made of bilayer and bolalipid molecules. (A) Onset energy required to form the membrane bud, 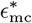, as function of bilayer head fraction 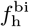 (for the parameters defined in Fig. 3.) (B and C) Fraction of U-shaped bolalipid molecules *u*_f_ (B) and bilayer molecules (C) in the outer and inner layers of the membrane bud and in the flat mother membrane as a function of the bilayer head fraction 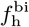. Top panels show the respective snapshots of membrane surface around cargo, where bilayer lipids are shown in light blue as in Fig. 4. (D) Average diameter of transient pores in the membrane bud and the mother membrane as function of bilayer head fraction 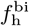 and respective snapshots of membrane leaflet surfaces surrounding the bud (top panel).

Importantly, we observed nearly no pores in the membrane bud in mixed membranes, even when we only had a very small fraction of bilayer-forming lipids (Fig. 5D). Only as the bilayer fraction increased, did we observe the formation of very small pores in the bud. For the flat mother membrane, however, membrane pores started to form with increasing values of *f* ^bi^. They acquired sizes similar to those obtained around the bud in pure bolalipid membranes (Fig. 4D). The pore formation in the flat mother membrane is likely promoted because the membrane becomes destabilized by the increasing proportion of bilayer lipids which are close to the gas phase. Taken together, bolalipids can bud porelessly when bilayer-forming lipids, which cause membrane softening, are included.

## Discussion

All biological cells are enclosed in fluid lipid membranes. These must be both sturdy and flexible to contain the cellular interior and allow membrane reshaping during division and vesicle formation. Across nature, mono- and bilayer membranes, composed of bolalipids and bilayer lipids, have evolved to fulfil these partially conflicting requirements. In this work, we have developed the first minimal model that allows us to transparently compare the distinct behaviour of bilayer and bolalipid membranes. In our model, bolalipids are formed by joining together two bilayer lipids, with an adjustable molecular stiffness at the hinge point. Using our model we find striking differences between bilayer and bolalipid membranes in terms of thermal and mechanical stability, bending rigidity and presence of hypoelasticity. Our results highlight how bolalipid membrane behave effectively as two-component systems, exchanging bolalipids between different conformations and thereby allowing the material properties to be adapted to support membrane reshaping.

While flexible bolalipid membranes are liquid under the same conditions as bilayer membranes, we found that stiff bolalipids form membranes that operate in the liquid regime at higher temperatures. These results agree well with previous molecular dynamics simulations that suggested that bolalipid membranes are more ordered and have a reduced diffusivity compared to bilayer membranes [24, 29]. In our simulations, this is due to the fact that completely flexible bolalipids molecules adopt both straight (transmembrane) as well as the U-shaped (loop) conformation with approximately the same frequency. In contrast, stiff bolalipids typically only take on the straight conformation when assembled in a membrane. These results agree with the previous coarse-grained molecular dynamics simulations using the MARTINI force field which showed that the ratio of straight to U-shaped bolalipids increased upon stiffening the linker between the lipid tails [29].

While previous coarse-grained simulations predicted that bolalipids spontaneously transition between the straight and U-shaped conformations [29], how this happens in archaeal membranes and whether membrane proteins are involved in this conformational transition needs to be clarified in the future. Experimental studies suggest that archaeal membranes contain flippases and scramblases for the transitioning of bilayer lipids between membrane leaflets [43, 44], raising the possibility that similar proteins could also facilitate conformational transitions in bolalipids. In addition, it has been suggested that the viral fusion protein hemagglutinin could cause a transition from straight to U-shaped bolalipid conformation during the fusion of bolalipid vesicles with influenza viruses [17]. However, future investigation is required.

When we determined the bending rigidity of bolalipid membranes by measuring their response to thermal fluctuations, we found that membranes made from flexible bolalipids are only slightly more rigid than bilayer membranes. This result is consistent with previous atomistic simulations, which showed that the membrane rigidity was similar for membranes composed of bilayer lipids and flexible synthetic bolalipids [45]. Moreover, the result is consistent with a continuum theory which predicted that the rigidity of membranes formed of triblock copolymers is 20% larger than that of diblock copolymers [46]. However, bolalipids in extremophilic archaea are not predicted to be fully flexible as they are expected to pack tighter due to a large number of cyclopentane rings in the lipid tails [7, 33]. Indeed, we found that membranes made of stiff bolalipid molecules can exhibit stiffness that is more than an order of magnitude larger than that of bilayer lipids at the same membrane fluidity.

It is striking that membranes made from stiffer bolalipids showed a curvature-dependent bending modulus, which is a clear signature that bolalipid membranes exhibit hypoelastic behaviour during membrane reshaping. Another marked difference between bilayer and flexible bolalipid membranes is the ratio of the Gaussian rigidity to the bending modulus. Instead of being around −1 as for bilayer membranes [41], it is around −1/2 and therefore only half of that of bilayer lipids. It is not obvious how the Gaussian bending modulus would behave upon increasing bolalipid stiffness (*k*_bola_ *>* 0), or how to measure it due to the coupling between curvature and rigidity in bolalipid membranes. Membrane remodelling, such as the fission of one spherical vesicle into two, increases the bending energy by 8*πκ* but decreases the energy related to the Gaussian modulus by 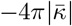[39], giving rise to a fission energy barrier of 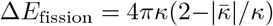. Our results indicated that while in bolalipid membranes *κ* is larger, 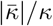 is smaller compared to bilayer membranes. Our results thus predict a larger energy barrier for membrane fission Δ*E*_fission_ in bolalipid membranes compared to bilayer membranes. It is tempting to speculate that this is one of the reasons why eukaryotes use bilayer membranes, enabling dynamic membrane remodelling and trafficking.

It is interesting to draw a parallel between monolayer membranes made of stiff bolalipid molecules and macroscopic membranes composed of rigid straight colloidal particles, which are geometrically similar, but living at different scales [47]. It has been found that colloidal membranes at these macroscopic scales follow the standard Helfrich theory for bilayer membranes [48], with rigidity that is three orders of magnitude higher than those of lipid bilayers [49]. In this case the tilt modulus was not pertinent, likely due to macroscopic system sizes. Similarly, we expect that the bending rigidity can be determined from membrane fluctuations independently of the tilt modulus for bolalipid membranes if they are prepared at similar relative sizes. Beyond quantitative differences, the comparison shows that monolayer membranes follow the same physics across many orders of magnitudes. However, when considering subcellular scales and the formation of high curvature, as in vesicle budding, the tilt modulus and thus the substructure of the membrane is expected to matter.

Our model makes a number of predictions that could be tested by experiment either in cells or in vitro. First, it predicts that a small increase in the fraction of archaeal bilayer lipids should be sufficient to soften a bolalipid-rich membrane. While this could be tested in the future, so far only very few studies have yet reported experimental analysis of archaeal membrane mixtures [18, 50]. Second, we observed that membranes with moderate bolalipid molecular rigidity *k*_bola_ exhibit curvature-dependent bending rigidity. To experimentally verify this, one could extrude membrane tethers from cells while controlling for membrane tension. Finally, to get to the core mechanism underlying our findings, it will be important to develop experimental methods that will allow the fraction of U-shaped bolalipid conformers per leaflet to be imaged and measured.

We found that membranes formed of a mixture of bilayer and bolalipids, similar to archaeal membranes, function as a composite liquid membrane that softens when adding bilayer lipids. However, while in our simulations the bending rigidity monotonically decreases with bilayer fraction, previous experiments of mixture membranes of bilayer and bolalipids with cyclopentane rings suggested that the bending rigidity non-monotonically depends on the fraction of the membrane made up of bilayer lipids [18]. It remains to be determined whether the result is due to the specific lipids used, the resulting mismatch in lengths between two stacked bilayer lipids and a straight bolalipid, the experimental conditions or to non-linear effects such as the formation of lipid domains that soften the membrane with increasing bolalipid content. The same experiments reported that membranes consisting solely of bolalipids are more rigid than bilayer membranes and non-fluctuating, which is in agreement with our high bending modulus simulation results for near-pure bolalipid mixture membranes.

To investigate how bolalipid membranes respond to changing membrane curvature, we performed simulations in which small cargo particles budded from flat membranes. We found that by enforcing curvature on bolalipid membranes, the fraction of U-shaped bolalipids increased around the cargo bud, especially in the outer membrane layer and hence softened the membrane. As another mechanism to release curvature stress we observed the formation of membrane pores, which could be mended by adding small amounts of bilayer lipids, similar to the mixture membranes that are found in archaea [8]. Our results suggest that enforcing membrane bending can soften bolalipid membranes locally by increasing the number of U-shapes, rendering the membrane a mechanically switchable material where large curvature decreases stiffness.

Taken together, our results show how membranes which are mixtures of bilayer and bolalipids maintain cell integrity at high temperatures, while also undergoing leak free membrane bending. This suggest that archaeal membranes can balance opposing needs when adapting to extreme environmental conditions. Beyond understanding membrane properties and reshaping across the tree of life, these results pave the way for synthetic bolalipid membranes and bolalipid membrane containers with new and exciting material properties.

## AUTHOR CONTRIBUTIONS

MA performed computer simulations and analysis and wrote the manuscript. FF performed patch closing simulations, contributed to computer simulations, performed theoretical analysis, and wrote the manuscript. XJ contributed to budding simulations and the manuscript. BB co-supervised the project and edited the manuscript. AŠ conceived the study, guided the project, and wrote the manuscript.

## DATA AVAILABILITY

The simulation input files and codes are freely available at [51].

## ACKNOWLEDGEMENTS

MA, BB, and AŠ acknowledge funding by the Volkswagen Foundation Grant Az 96727. FF acknowledges financial support by the NOMIS foundation. AŠ acknowledges funding by ERC Starting Grant “NEPA” 802960. We thank Claudia Flandoli for help with illustrations.

