## Supplemental Information for "Stability vs flexibility: reshaping archaeal membranes in silico"

### 1. MEMBRANE MODEL

To study the topological difference between bilayer lipids and bolalipids, we started from a coarse-grained model that was previously developed for lipid bilayer membranes by Cooke and Deserno [1]. Each lipid is composed of a chain of beads. Bilayer lipids are composed of one head bead connected to a chain of 2 tail beads. In contrast, bolalipids are composed of two head beads linked to a chain of 4 tail beads (see Fig. 1C). Importantly, bilayer and bolalipids have identical head groups.

Each adjacent pair of beads in a lipid is connected by a finite extensible non-linear elastic bond (FENE). For a given bond length $r$, its potential is the sum of an attractive term and a purely repulsive Lennard-Jones potential that enforces volume exclusion

$$U_{\text{bond}}(r) = -\frac{1}{2} K R_0^2 \ln\left(1 - \left(\frac{r}{R_0}\right)^2\right) + U_{\text{lj}}(r), \quad r \in [0, R_0], \tag{S1}$$

with $K = 30 k_{\text{B}} T/\sigma^2$ and maximum length $R_0 = 1.5\sigma$ in the first term. We note that in the simulations our time, distance and energy units are respectively $\tau$, $\sigma$ and $k_{\text{B}} T$, and the Boltzmann constant $k_B = 1$. Consequently, our unit of mass is given by $1m = 1 k_{\text{B}} T \tau^2/\sigma^2$. The second term of Eq. (S1) is given by

$$U_{\text{lj}}(r) = U_{\text{m}} \cdot \left(x^{-12} - 2x^{-6} + 1\right), \tag{S2}$$

where $x = \min(r, r_{\text{c}})/r_m$, where $r_m$ is where the potential reaches its minimum value $U_{\text{m}}$ and $r_{\text{c}}$ is its cutoff. For bonded beads, we set $U_m = 1 k_{\text{B}} T$, $r_{\text{m}} = r_{\text{c}} = 2^{1/6}\sigma$, so that the repulsive part is zero for $r > r_{\text{c}}$.

For non-bonded beads, as a first term we pick also a purely repulsive form with $r_{\text{m}} = r_{\text{c}} = 2^{1/6}\sigma \approx 1.12\sigma$, and parametrize on its strength by scanning the interval $U_{\text{m}} = \epsilon_{\text{p}} \in [0.5, 2] k_{\text{B}} T$. The inverse of this potential depth is the effective temperature $T_{\text{eff}} = k_{\text{B}} T/\epsilon_{\text{p}}$ in our system. Following Cooke and Deserno [1], we scaled down $r_{\text{m}}, r_{\text{c}}$ by 0.95 for interactions with head beads, to ensure no spontaneous curvature in a membrane leaflet [1],[2]. Accordingly, we note that in our snapshots each head bead is represented as a sphere of diameter $0.95\sigma$ and each tail bead as a bead of diameter $1\sigma$.

To model lipid rigidity between each consecutive three beads $b_1, b_2, b_3$ in a lipid we set a harmonic angle potential

$$U_{\text{angle}}(\theta) = K \cdot (\theta - \pi)^2, \quad \theta \in [0, \pi], \tag{S3}$$

with $\theta = \widehat{b_1, b_2, b_3}$. If one of the beads is a head bead (i.e. in a bilayer lipid), we set $K = k_0 = 5\, k_{\text{B}} T$. In contrast, for the two inner angles in a bolalipid we set the interval $K = k_{\text{bola}}$, where $k_{\text{bola}}$ is a global constant in the simulation, equal for all bolalipids. In our work, for different simulations, we vary it in the interval $[0, 5] k_{\text{B}} T$. Therefore, the difference between a bolalipid and two bilayer-forming lipid molecules is the extra bond and two equal angle potentials keeping each three bead connected and aligned, respectively.

To model the (implicit) hydrophobic interaction between lipid tail beads, we add a longer range attractive cosine squared potential

$$U_{\mathrm{cs}}(r) = -\epsilon_{\mathrm{p}} \cos^2\left(\frac{\pi}{2}\mathrm{clip}\left(\frac{r-r_c}{\omega}, 0, 1\right)\right)$$
$$= \begin{cases} -\epsilon_{\mathrm{p}} & r \leq r_c \\ -\epsilon_{\mathrm{p}} \cos^2\left(\frac{\pi}{2}\frac{r-r_c}{\omega}\right) & r_c < r < r_c + \omega \\ 0 & r \geq r_c + \omega \end{cases} \tag{S4}$$

where $\mathrm{clip}(x, a, b) = \max(\min(x, b), a)$ and $r_c$ is set to the cutoff $2^{1/6}\sigma$ of the volume exclusion interaction. The repulsive Lennard-Jones and the attractive potential are combined to make the tail-tail interaction repulsive in the range $[0, r_c]$ and attractive in the range $[r_c, r_c + \omega]$. We take $\omega$, the attractive range width, as a parameter. As we joined two bilayer lipids to form a bolalipid, in our model bilayer lipids and bolalipids share the same hydrophobic interaction.

### 2. SIMULATION PROTOCOL FOR MEMBRANE SELF-ASSEMBLY

We initially verified self-assembly of our lipids into a flat membrane: we placed lipids in a dispersed gas configuration in a periodic 3D cube. We then evolved the system with timestep $\delta t = 0.01\,\tau$ under a Langevin thermostat with relaxation time of $1\tau$ and checked a flat membrane patch eventually formed (see Movie S1).

For the rest of our simulations we pre-assembled the membrane. This was done by placing lipids far from each other in a hexagonal grid, locking the lipids vertical position and conformation, compressing with volume exclusion until a target area per lipid is reached, followed by energy minimization and a final check that the lipids formed a horizontal single cluster.

### 3. BOLALIPID CONFORMATION IN FLAT MEMBRANES

For simulating flat membrane patches of bolalipids, we combined the previously used Langevin thermostat with relaxation time of $1\tau$ with a Nosé–Hoover barostat with relaxation time of $10\tau$. In LAMMPS this amounts to combining the commands 'fix langevin' with 'fix nph'. We configured the barostat to set lateral pressure $P_{\mathrm{xy}}$ to zero by re-scaling the simulation box in the $x$-$y$ plane. We compare this setup to a fixed box length setup, and a NPT ensemble setup, in SI section 17.

For a bolalipid in a flat membrane, we found thresholding on the angle between its first and second half $\theta = \angle\overrightarrow{b_3, b_1}, \overrightarrow{b_4, b_6}$ is sufficient to classify its conformation (see Fig. 1E, right). We checked for a few parameters that this angle follows a bimodal distribution (Fig. S1) and that indeed bolalipids assume either a straight conformation, one head bead in opposing membrane leaflets, with $\theta \approx \pi$ or a U-shaped conformation, both head beads on the same leaflet, with $\theta \approx 0$. Accordingly, we marked a bolalipid as being in the U-shape if $\theta < \pi/2$ and in the straight conformation otherwise (see Fig. 1E for a snapshot).

For pre-assembly of flat membranes containing bolalipids, this implied a choice of which conformation to pick for each lipid, straight or U-shaped. We verified that for the limit cases of flexible and rigid bolalipids, starting the system with all bolalipids in the straight and U conformation, respectively, resulted in the U-shaped bolalipid fraction $u_{\mathrm{f}}$ equilibrating after simulating at most for $10^4\tau$ (Fig. S2).

After verifying this, we started our simulations with a fraction of bolalipids derived from assuming a two-state system, with $\Delta E$ given solely by the central angle potentials (e.g., for U-shaped bolalipids both angles assume values of $\pi/2$). We checked our system had equilibrated quantitatively by

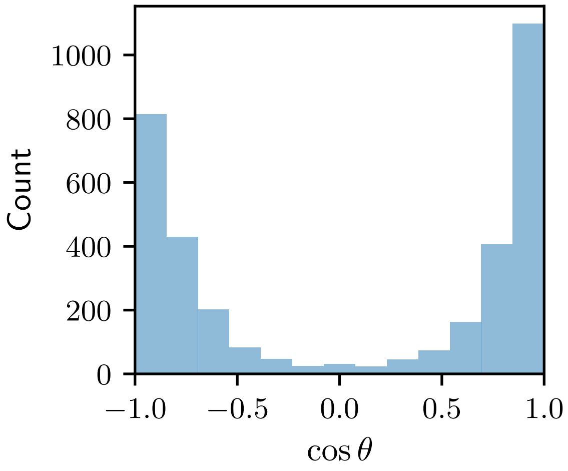


Figure S1. Histogram of the cosine of the angle $\theta$ between halves of a bolalipid ($k_{\mathrm{bola}} = 0$) for the last frame of the simulation of a flat membrane of flexible bolalipids.

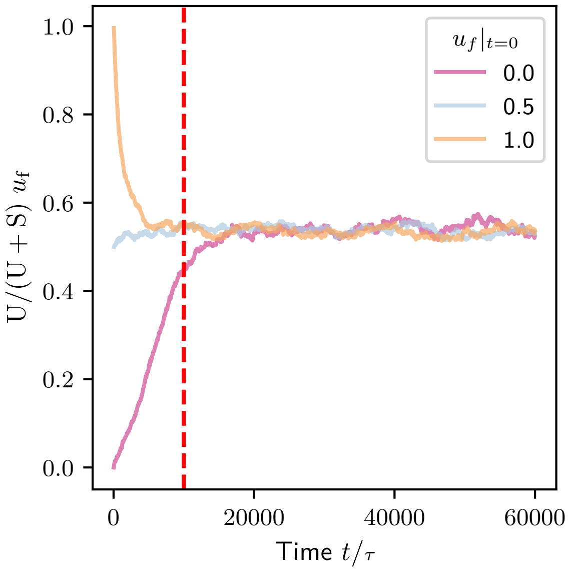


Figure S2. Time series of $u_f$, the fraction of bolalipids in the U-shaped conformation, for the simulations done for flexible bolalipids ($k_{\mathrm{bola}} = 0$), where the system was pre-assembled with three different initial values $u_f|_{t=0}$. The red dashed line marks the typical equilibration time of $10^4\tau$.

observing that the time series of the fraction of bolalipids in the U-shaped conformation approached a steady state (Fig. S3). We still equilibrated for $10^4\tau$ before taking measures.

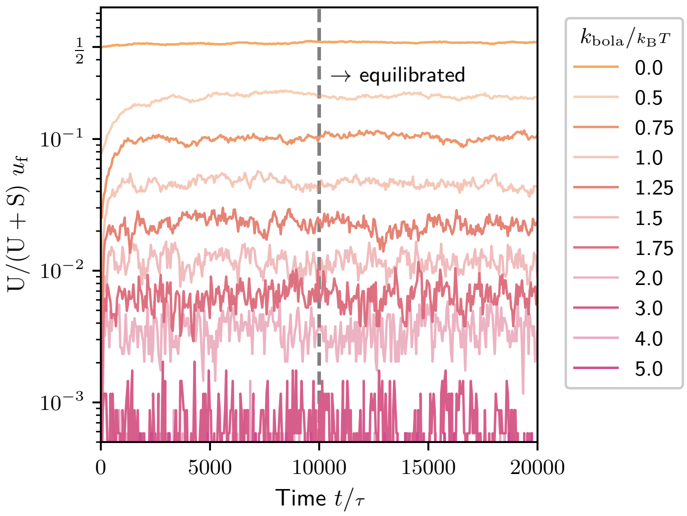


Figure S3. Time series of fraction of bolalipids in the U-shaped conformation $u_f$, for simulations of pure bolalipid membranes with different values of the bolalipid rigidity $k_{\mathrm{bola}}$. The dashed line marks the timestamp used for defining the system as equilibrated. All equilibrium measures are taken from frames after this point.

### 4. SIMULATION PROTOCOL FOR MEMBRANE PHASE DETERMINATION

For phase determination (Fig. 1F), the membrane was assembled in a horizontal grid of lipids with approximately $25^2$ head beads in each leaflet. For bolalipid membranes, we set the initial fraction of U-shaped bolalipids as we did for simulations depicted in Fig. S3. We then ran the simulation, halting it automatically if the box size diverged, at which point we could consider the membrane as being in the gas phase. Otherwise, the simulation ran to completion for $\Delta_{\mathrm{total}} = 10 \times 10^3\,\tau$. This was determined to be sufficient for both the box size $L$ and the relative amounts of lipid species and conformations to equilibrate.

### 5. DIFFUSION CONSTANT

For computing the diffusion constant $D$, we first found the corresponding mean square displacement for each $\Delta t$ and then fitted $D = \left\langle |\Delta x|^2 \right\rangle / (\Delta t)$. To compute the M.S.D. of a specific lipid in a temporary conformation, we averaged $|\Delta x|$ over all possible intervals of duration $\Delta t$ where the conformation was held. We took care to exclude displacements of lipids floating in the gas phase of the simulation. Fig. S4 shows $D$ as a function of temperature $T_{\mathrm{eff}} = k_{\mathrm{B}}T/\epsilon_{\mathrm{p}}$ for the different membrane types. Fig. S4 shows a discontinuity of the diffusion constant $D$ as a function of $T_{\mathrm{eff}}$. We took this discontinuity as marking the transition of the membrane from the gel phase to the liquid phase, and determined it for bilayer and bolalipid membranes for different values of $k_{\mathrm{bola}}$ (Fig. S4). For all membranes and parameters tested this is equivalent to setting the minimum diffusion constant for a liquid membrane to $D = 5 \times 10^{-4}\,\sigma^2/\tau$.

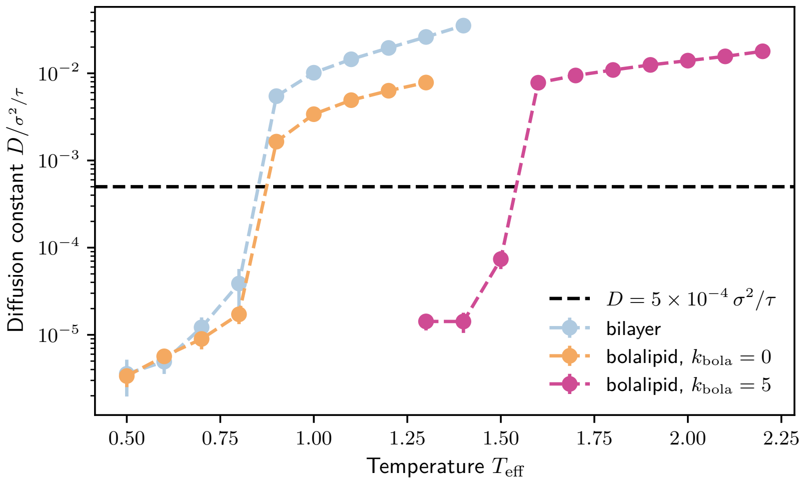


Figure S4. Diffusion constant for different pure membranes at $w = 1.5\,\sigma$ versus temperature $T_{\mathrm{eff}}$. The discontinuity (jump) in $D$ marks the transition from the gel phase to the liquid phase. The dashed black line marks the minimum diffusion constant that is required to classify as a liquid membrane at $D = 5\times10^{-4}\,\sigma^2/\tau$.

### 6. TWO-STATE MODEL FOR BOLALIPID CONFORMATION

We judged our two-state model fitness in Fig. 2B by first making the model linear, expressing it as $\log(1/u_{\mathrm{f}} - 1) = x_1 k_{\mathrm{bola}} + x_0$; then we restricted it to points where $k_{\mathrm{bola}} \leq 2\,k_{\mathrm{B}}T$, or, equivalently, when there were on average $\approx 2$ or more U-shaped bolalipids; with this choices we obtained $R^2 = 0.99$.

By splitting the average potential energy between an internal contribution (bonds, angles and pair interactions between particles in the same molecule) and an external contribution (pair interactions between a molecule and its neighbours), we determined the transition energy from straight to U-shaped bolalipids in detail. We found that this transition lowers the internal potential energy of the bolalipid while increasing its interaction energy. In total, we obtained an energy barrier for the transition of $\Delta E_{\mathrm{s}\to\mathrm{u}} = 0.79 \pm 0.01\,k_{\mathrm{B}}T$. Since the fit indicates, however, that the U-shaped bolalipid conformation is preferred over the straight conformation, we conclude that there must be either an entropic contribution to the free energy or an intermolecular interaction energy favouring U-shaped bolalipids.

### 7. FLUCTUATION SPECTRUM

In order to measure height fluctuation spectrums (Fig. 2C, and Fig. 3B), we used membranes with $60^2$ head beads per leaflet, with minimum $\Delta_{\mathrm{eq}}$ set to $20\times10^3\tau$; total runtime was $\Delta_{\mathrm{total}} = 60\times10^3\,\tau$.

For measuring the bending modulus $\kappa$, we follow the analysis done by Cooke and Deserno [1]. We simulate a horizontal membrane in a periodic simulation box of dimensions $(l_{\mathrm{x}}, l_{\mathrm{y}}, l_{\mathrm{z}})$, that is horizontally square with $l_{\mathrm{x}} = l_{\mathrm{y}} = L$. Importantly, we keep membrane tension to a minimum by setting the lateral pressure $P_{\mathrm{xx}} = P_{\mathrm{yy}} = 0$ via a barostat. As a first equilibration check, we consider the lateral box size $L$ time series. Starting by considering the full series, we measure how much the first and last half differ. For each half, we compute the maximum, minimum and the diameter (max - min). If the relative difference is less than 30%, we consider it equilibrated. Otherwise, we exclude

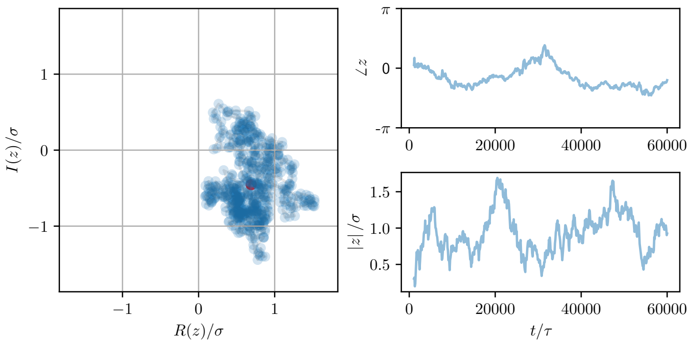


(A) non-equilibrated mode at $n = (1, 0)$.

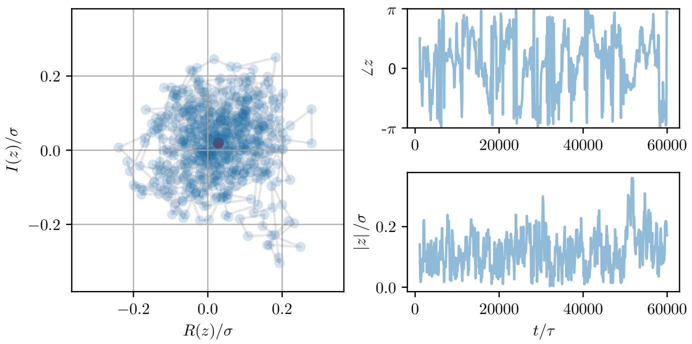


(B) equilibrated mode at $n = (2, 0)$.

Figure S5. Path of the complex component of the fourier transform of membrane height at $n = (1, 0)$ (A) and $n = (2, 0)$ (B) for bolalipid pure membrane at $k_{\mathrm{bola}} = 0.5k_{\mathrm{B}}T$. Left plots show the trajectory in the complex plane, while on the right we plot their phase and norm versus time. The mode in (A), with an autocorrelation time of roughly $10^4\tau$, has only 10 uncorrelated points. On the other hand the mode in (B), with an autocorrelation time of approximately $10^3\tau$, has $\approx 60$ uncorrelated samples and thus crosses the chosen threshold of 20 samples for being considered equilibrated.

the first frame of the time series, and repeat the check.

For each of the $N_{\mathrm{f}}$ remaining frames, we intend to obtain the height field of the membrane $h(x, y)$ within the $x$-$y$ plane. However, our simulations are particle-based and hence our system is discrete. Therefore, we divide the horizontal plane in a regular $m \times m$ grid, where $m$ is the length of a grid cell. We set $m = 40$ by trial and error so that no bins will be empty. We compute $h_b$, the average height of bin $b$. We then apply the 2D discrete Fourier transform in the $x$-$y$ plane, obtaining complex components $h_n$ for $n \in [-m/2, ..., 0, .., m/2] \times [-m/2, ..., 0, .., m/2]$, with $h_n = h_{-n}$.

We correct for the binning by multiplying by $\mathrm{sinc}(n_x/m)\,\mathrm{sinc}(n_y/n)$, which mostly affects the smallest wavelengths [1]; we also scale by $\langle L\rangle$ obtaining $u = h\langle L\rangle$. For each mode number vector $n = (n_x, n_y)$ of $u$, we compute its amplitude squared $|u_n|^2$ and phase $\angle u_n$ (examples in Fig. S5). For both the amplitude and phase we then compute the autocorrelation time $\tau_c$ in order to get the statistical inefficiency $g = 2\tau_c + 1$. The phase should regularly jump through the endpoints $[0, \pi]$, so checking it too for stationarity allows to exclude modes whose amplitude is static but which have their phase stuck. By doing this, we find that we can improve the analysis compared to the current state of the art [3], which only checks the amplitude squared. Between both the amplitude squared and phase components, we take the largest $g$, which then gives us the number of uncorrelated data points as $N_\mathrm{f}/g$. We accept a mode as equilibrated if the remaining trajectory contains at least 20 uncorrelated data points. The standard deviation of the mean of $|u_n|^2$ must then be scaled by $\sqrt{g}$.

For each spectrum measurement, we performed four simulations with different seeds for the thermal noise. We retained only modes which had equilibrated on all replicas and averaged over modes with the same mode number. We checked that the box size $L$ varied less than 1% between replicas. To not impose an artificial variable window in mode number, we computed the first equilibrated mode for all our simulations and took the maximum equilibrated mode number as a global minimum threshold. We found in our simulations $n \geq 2$. This roughly corresponds to a maximum wavelength cutoff of $32\sigma$ given that our simulation boxes have length $60\sigma$. We used twice the membrane thickness, i.e., $12\sigma$ as minimum wavelength cutoff.

According to continuum membrane theory, at zero membrane tension the fluctuation spectrum is given by [4]

$$\langle |h_n|^2 \rangle = \frac{k_\mathrm{B}T}{L^2 \kappa q^4}\,, \tag{S5}$$

where $\kappa$ is the bending modulus, $L$ is the box size, $q = 2\pi n/L$ is the (angular) wave number and $n$ is the mode number[1]. While this was adequate for bilayer membranes, we needed to add the tilt term, parametrized by the tilt modulus $\kappa_\theta$ (see main text Eq. (1), [5]) to get reasonable fits for rigid bolalipid membranes. Then, we obtained

$$\langle |h_n|^2 \rangle = \frac{k_\mathrm{B}T}{L^2} \left( \frac{1}{\kappa q^4} + \frac{1}{\kappa_\theta q^2} \right)\,. \tag{S6}$$

We fitted the data using $N$ measurements of mean and mean standard deviation $(y_i, \sigma_i, x_i)$, with $y = \langle |u_n|^2 \rangle = \langle L \rangle^2 \langle |h_n|^2 \rangle$ and $x = q = 2\pi |n| / \langle L \rangle$ to each model $f(x_i) = y_i$. In this case, the different models $f(x)$ are given by Eqs. (S5) and (S6), where Eq. (S5) can be derived from Eq. (S6) by formally setting $\kappa_\theta = \infty$. Typical example fits are shown in Fig. S6.

We used the reduced $R^2$ value as an indicator of goodness of fit, defined as $R^2 = \sum_i ((f(x_i) - y_i)/\sigma_i)^2/N$. Reasonable values were recognized as $R^2 \leq 1$. $R^2$ is plotted together with fit results for pure bolalipid membranes and bolalipid/bilayer mixture membranes in Fig. S7. In the first row we show the results of the fit of Eq. (S5) while in the second row we used Eq. (S6). In general, for small values of bolalipid rigidity or large bilayer fraction the fits without the tilt term were still reasonable. However, as the bolalipid rigidity increased or the bilayer fraction decreased the fits became worse. Then only fits with the tilt term were reasonable. In addition, we plot the same measures for temperature sweeps for the bilayer at $T_\mathrm{eff} = 1.2$, the flexible bolalipid and the rigid bolalipid membranes in Fig. S8. The results are summarized for all membrane types in Fig. S9.

We omitted the error in the main text when less than the unit. Moreover, the low values of tilt modulus will necessarily be accompanied of smaller error bars since the smaller the value, the larger the influence the term will have on the amplitudes.

We note that in our large flat membrane simulations ($L \approx 60\sigma$), flexible bolalipid membranes at $\omega = 1.5\sigma$ are stable only at temperatures smaller than $T_\mathrm{eff} < 1.4$. For $T_\mathrm{eff} = 1.4$ and presumably above, the membrane folds while shrinking the box until self-contact occurs. This is accompanied by massive oscillations of the box pressure, and can be avoided by halving the timestep. Since these points are near the gas phase and halving the timestep did not significantly change measurements for non-collapsing membranes (see SI section 17), we simply kept the timestep at $0.01\,\tau$. On the other hand, both the bilayer membranes and rigid bolalipid membranes of same size disassemble by pore formation, followed by simulation box expansion in response to the increased pressure, respectively at $T_\mathrm{eff} = 1.5$ and 2.3. This explains why in Fig. S9 we have data for bilayer at higher $T_\mathrm{eff}$ than for the flexible bolalipids.

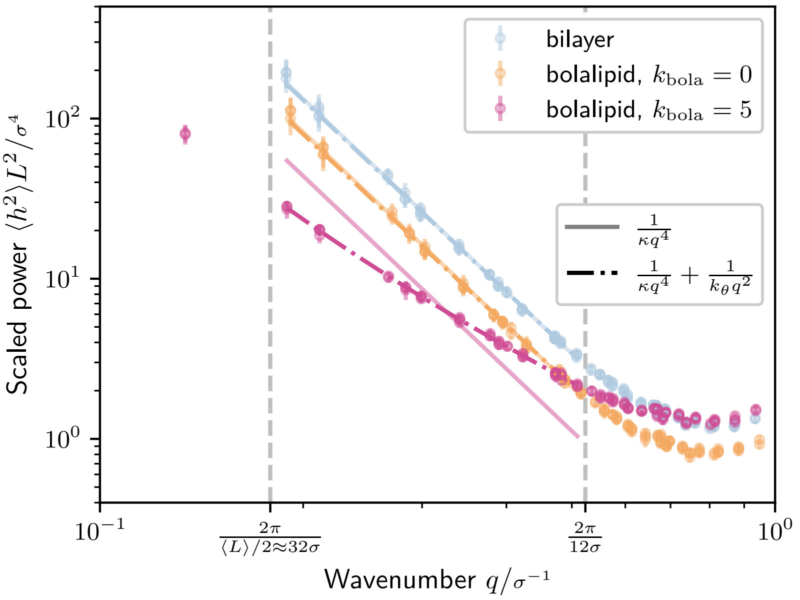


Figure S6. Fluctuation spectra and corresponding fits according to Eq. (S6) for a model without and with tilt modulus, for a bilayer membrane at $T_{\rm eff} = 1.2$, a flexible bolalipid membrane ($k_{\rm bola} = 0$) and a rigid bolalipid membrane ($k_{\rm bola} = 5\, k_{\rm B}T$). In vertical dashed lines we marked the interval of wave numbers that selects the modes used for fitting. The first rigid bolalipid equilibrated mode is to the left of this interval and thus excluded from the corresponding fit.

We also note that for both the bilayer and the flexible bolalipid membrane, in $T_{\rm eff} \in [1.2, 1.3]$, the tilt modulus varies non-monotonically. It first increases, aligning with the expectation that higher pair potential temperature would reduce order and thus increase the cost of local coordinated tilting, i.e., increasing the tilt modulus $\kappa_\theta$. However, at the higher $T_{\rm eff}$ it abnormally decreases.

Comparing the bending moduli of the flexible bolalipid membrane ($\kappa \approx 8\, k_{\rm B}T$, $\kappa_\theta \approx 30 \pm 10\, k_{\rm B}T/\sigma^2$) to the bilayer membrane ($\kappa \approx 5\, k_{\rm B}T$, $\kappa_\theta \approx 33 \pm 20\, k_{\rm B}T/\sigma^2$, cf. Fig. S7B) we find similar results. By systematically investigating the membrane rigidity as a function of temperature, we found that in general flexible bolalipid membranes have a slightly increased rigidity compared to bilayer membranes (Fig. S9). We also verified that by increasing $T_{\rm eff}$, a rigid bolalipid membrane softens in the same manner as bilayer and flexible bolalipid membranes (Fig. S9).

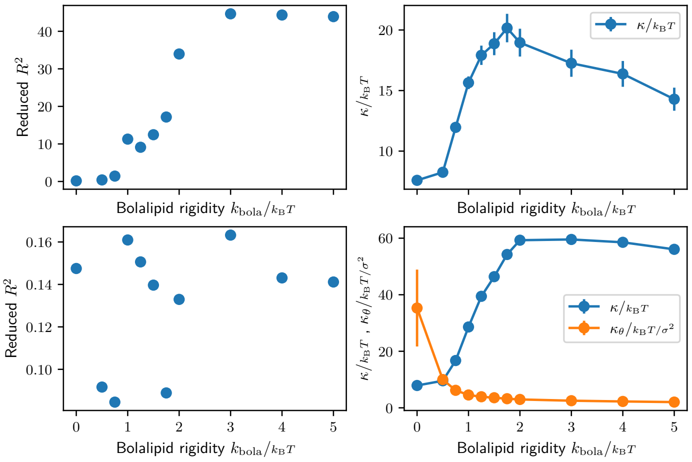


(A) Bolalipid Membranes

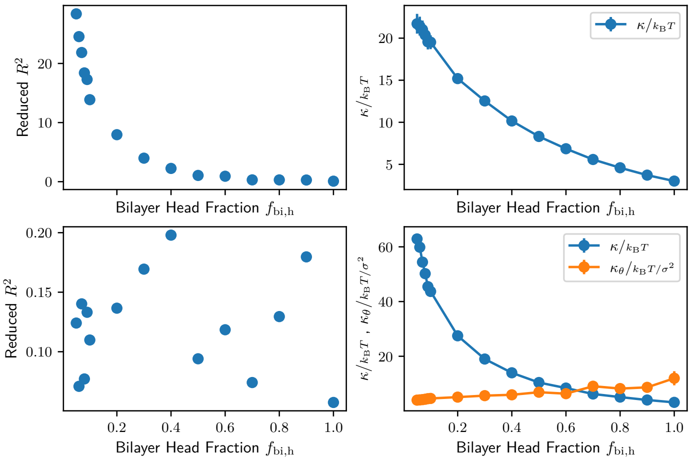


(B) Mixture Membranes

Figure S7. Fluctuation spectrum fit comparisons for pure bolalipid membranes (A) and bolalipid/bilayer mixture membranes (B). In the first row we show the results of the fit of Eq. (S5) while in the second row we used Eq. (S6).

### 8. MEASURING BENDING MODULUS VIA MEMBRANE TETHER/CYLINDER SIMULATIONS

Since the method is described elsewhere [6], here we detail specifics. For all measurements we ran 4 seeds, each over $20 \times 10^3\,\tau$, integrating in the NVT ensemble using a Langevin thermostat with damping coefficient $1\tau$ and unit temperature. The membrane was assembled into a cylindrical shape, with the number of heads in each leaflet pre-balanced. The stress tensor was measured at $1\tau$ intervals. The radius was measured from trajectory frames saved every $20\tau$ in the following way. First we excluded lipids in gas phase by clustering. Then we computed the centre of mass of the membrane and set it as our origin for the cylinder cross-section $x$-$y$ plane. Then we computed the

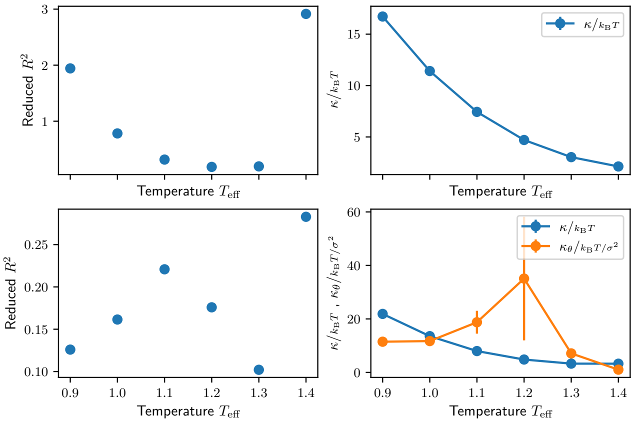


(A) Bilayer membranes fluctuation spectrum fits.

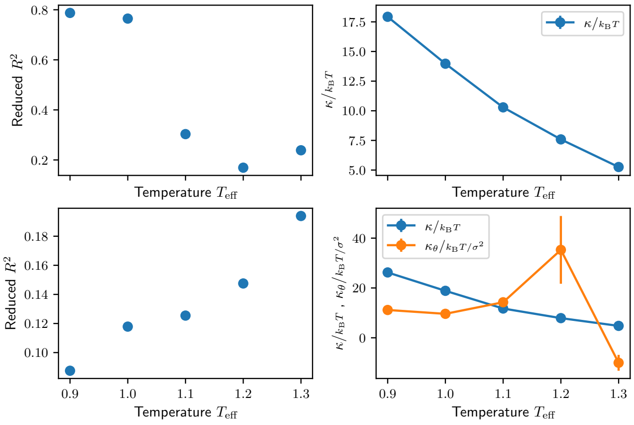


(B) Flexible bolalipid membranes fluctuation spectrum fits.

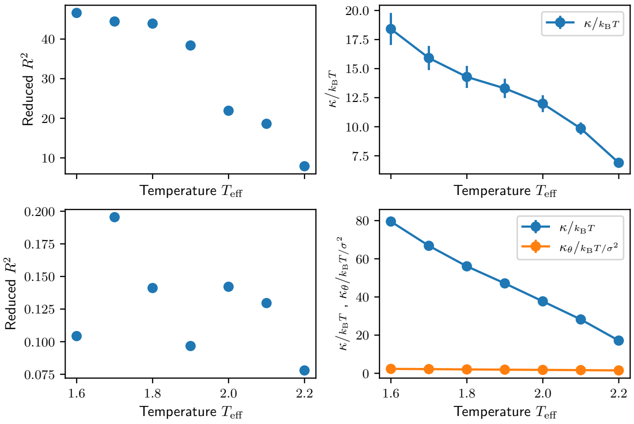


(C) Rigid bolalipid membranes fluctuation spectrum fits.

Figure S8. Fluctuation spectrum fit comparisons for pure bilayer membranes (A), flexible (B) and rigid pure bolalipid membranes (C). For each type of membrane, in the first row we show the results of the fit of Eq. (S5) while in the second row we used Eq. (S6), as a function of $T_{\rm eff}$.

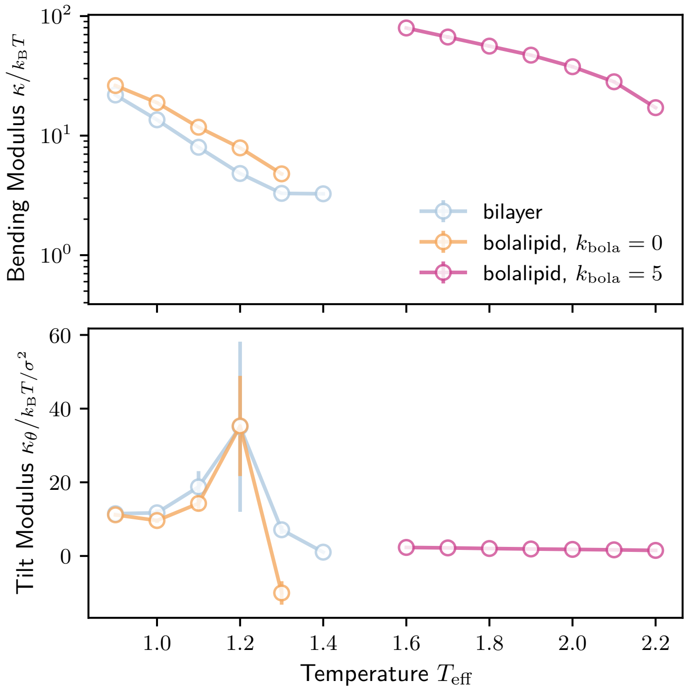


Figure S9. Fluctuation spectrum fit results, with (top) bending and (bottom) tilt modulus, for bilayer, flexible bolalipid and rigid bolalipid membranes at $w = 1.5$ as a function of $T_\mathrm{eff}$.

| $\Delta t$ | $N$ | $l_z$ | $H$ | $\left\lvert\frac{P_x+P_y}{2P_z}\right\rvert$ | $\kappa$ |
|---|---|---|---|---|---|
| 0.001 | 19457 | 120 | (3.08+/-0.04)e-02 | 0.03+/-0.06 | 8.9+/-2.4 |
| | 10000 | 70 | (3.447+/-0.004)e-02 | 0.03+/-0.05 | 8.4+/-1.1 |
| | | 80 | (3.923+/-0.008)e-02 | 0.04+/-0.04 | 8.1+/-0.6 |
| | 5000 | 40 | (3.948+/-0.006)e-02 | 0.05+/-0.05 | 8.8+/-1.3 |
| | | 50 | (4.942+/-0.009)e-02 | 0.011+/-0.021 | 8.6+/-0.6 |
| | | 60 | (5.857+/-0.005)e-02 | 0.018+/-0.017 | 8.22+/-0.33 |
| | | 70 | (6.82+/-0.02)e-02 | 0.010+/-0.013 | 8.27+/-0.30 |
| | | 80 | (7.17+/-0.03)e-02 | 0.009+/-0.011 | 8.7+/-0.3 |
| 0.005 | 5000 | 90 | (7.86+/-0.03)e-02 | 0.010+/-0.006 | 9.16+/-0.21 |
| | | 96 | (8.13+/-0.03)e-02 | 0.085+/-0.013 | 8.87+/-0.27 |

Table S1. Parameters and measurements for bilayer lipid cylinders. $N$ is number of lipids.

measured radius as $R = 1/\langle 1/r \rangle$, where the average is over beads and where $r = |x^2 + y^2|$ is the distance to the cylinder axial radius. Using $1/r$ instead of $r$ directly compensates to first order for the shell volume $2\pi r dr l_z$ dependency on $r$.

After qualitatively checking for quick equilibration of the radius, fraction of U-shaped conformers and the stress tensor components, we dispensed with lengthy equilibration, discarding only the first $1000\tau$ of the trajectory. These observables were then averaged over the rest of the trajectory, with errors determined in a blocking-equivalent manner.

Specifically, in these simulations we focused on three pairs of parameters, which we name bi-

| $\Delta t$ | $N$ | $l_z$ | $H$ | $\left\lvert\frac{P_x+P_y}{2P_z}\right\rvert$ | $\kappa$ | $u_f$ |
|---|---|---|---|---|---|---|
| | 19457 | 120 | (3.024+/-0.004)e-02 | 0.06+/-0.08 | 9.2+/-1.6 | 0.534+/-0.006 |
| | 10000 | 70 | (3.354+/-0.004)e-02 | 0.03+/-0.08 | 9.7+/-2.1 | 0.536+/-0.004 |
| | | 80 | (3.848+/-0.004)e-02 | 0.04+/-0.05 | 9.5+/-1.4 | 0.537+/-0.005 |
| 0.001 | 5000 | 40 | (3.907+/-0.006)e-02 | 0.04+/-0.07 | 9.6+/-2.0 | 0.531+/-0.004 |
| | | 50 | (4.821+/-0.004)e-02 | 0.03+/-0.04 | 9.7+/-1.0 | 0.542+/-0.005 |
| | | 60 | (5.672+/-0.006)e-02 | 0.021+/-0.024 | 9.3+/-0.6 | 0.544+/-0.005 |
| | | 70 | (6.35+/-0.02)e-02 | 0.020+/-0.018 | 9.2+/-0.5 | 0.546+/-0.005 |
| | | 80 | (7.19+/-0.03)e-02 | 0.008+/-0.012 | 9.1+/-0.4 | 0.551+/-0.005 |
| 0.005 | 5000 | 90 | (7.82+/-0.03)e-02 | 0.095+/-0.007 | 9.70+/-0.16 | 0.5583+/-0.0015 |
| | | 96 | (8.13+/-0.03)e-02 | 0.088+/-0.012 | 9.35+/-0.28 | 0.563+/-0.004 |

Table S2. Parameters and measurements for flexible bolalipids tethers.

| $\Delta t$ | $N$ | $l_z$ | $H$ | $\left\lvert\frac{P_x+P_y}{2P_z}\right\rvert$ | $\kappa$ | $u_f$ |
|---|---|---|---|---|---|---|
| 0.001 | 19457 | 120 | (2.921+/-0.006)e-02 | 0.020+/-0.032 | 23.3+/-2.1 | 0.07401+/-0.00033 |
| | 10000 | 70 | (3.260+/-0.002)e-02 | 0.043+/-0.028 | 23.3+/-1.7 | 0.0806+/-0.0009 |
| | | 80 | (3.743+/-0.002)e-02 | 0.021+/-0.022 | 22.0+/-1.5 | 0.0898+/-0.0009 |
| | 5000 | 50 | (4.526+/-0.003)e-02 | 0.009+/-0.022 | 19.5+/-1.0 | 0.1053+/-0.0010 |
| | | 60 | (5.375+/-0.004)e-02 | 0.010+/-0.015 | 17.8+/-0.5 | 0.1225+/-0.0008 |

Table S3. Parameters and measurements for bolalipid tethers at $k_{\mathrm{bola}} = 1\,k_{\mathrm{B}}T$.

layer ($T_{\mathrm{eff}} = 1.1$), flexible bolalipids ($k_{\mathrm{bola}} = 0, T_{\mathrm{eff}} = 1.2$), and slightly rigid bolalipids at $k_{\mathrm{bola}} = 1\,k_{\mathrm{B}}T$ ($T_{\mathrm{eff}} = 1.4$). To guarantee correct integration, we lowered the timestep until the ratio $|(P_x + P_y)/(2P_z)|$ was less than 0.1. We also balanced increasing the cylinder length, to improve statistics on the stress tensor measurement, against computational performance and the occurrence of long-term deviations from cylindrical shape at high aspect ratio $R/l_z$. We include the exact parameters and data used in the plots in the main text in Tables S1 to S3.

We will now in detail explain the different behaviour of flexible versus stiffer bolalipid membranes

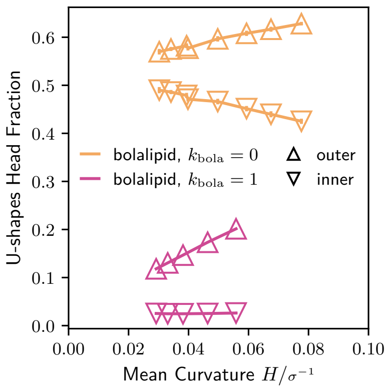


Figure S10. Fraction of lipid heads belonging to U-shaped bolalipids, for both outer (upwards triangles) and inner (downwards triangles) leaflets in cylinder membranes versus mean curvature $H$, for bolalipid membranes at $k_{\mathrm{bola}} = 0$ and $k_{\mathrm{bola}} = 1\,k_{\mathrm{B}}T$.

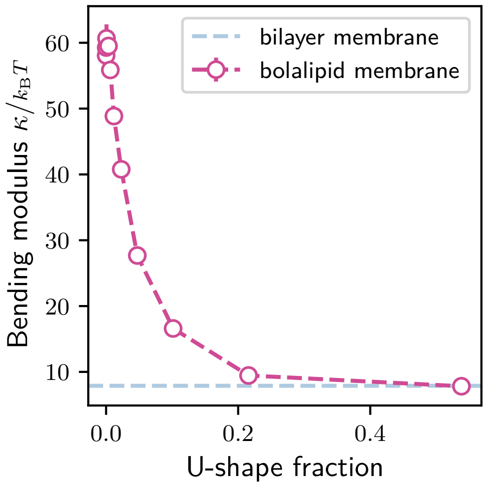


Figure S11. Bending modulus $\kappa$ versus U-shaped bolalipid fraction, using the same data as Figs. 2B and 2C. For reference the bilayer bending modulus at $T_{\text{eff}} = 1.1$ is shown in blue.

under bending. Membrane bending rigidity in bolalipid membranes decreases dramatically once a small fraction of U-shapes is allowed to form, but then plateaus once this U-shape fraction reaches 20%. In a curved bolalipid membrane, U-shapes must accumulate in the outer leaflet to accommodate for area difference. Together, the bending rigidity non-linear dependence on U-shape fraction, and the promotion of U-shapes by curvature, explain why in a membrane made of moderately stiff bolalipids ($k_{\text{bola}} = 1\,k_B T$), which contain very few U-shapes in the flat state, the bending rigidity of the membrane decreases as curvature increases. While in a membrane made of flexible bolalipid molecules ($k_{\text{bola}} = 0$), where many U-shapes are present in the flat membrane, the bending rigidity does not change with curvature.

Bending rigidity $\kappa$ in flat membranes composed of bolalipids decreases dramatically once a small fraction of U-shapes is allowed to form, but plateaus once more than 20% of U-shaped bolalipids are present. In details, our data shows that with an increasing bolalipid molecular rigidity $k_{\text{bola}}$, both the number of U-shaped bolalipids decreases (Fig. 2B) and the membrane rigidity $\kappa$ increases (Fig. 2C). Thus, the correlation suggests that U-shaped bolalipids soften the membrane, in a non-linear way where most of the change in membrane bending rigidity happens for U-shaped bolalipid fraction $< 20\%$ (Fig. S11).

Separately, membrane curvature affects the area difference between curved membrane leaflets and thus drives U-shape accumulation. To be specific, a cylindrical membrane with area $A$, mean curvature $H$ and thickness $h$ has the outer leaflet with area $A(1 + Hh)$ and the inner leaflet with smaller area $A(1-Hh)$. This can be large, in our simulations up to an area change of $Hh = 25\%$. For pure bolalipid membranes, straight bolalipids occupy the same space in each leaflet. Area difference can then be achieved only by having a different amount of U-shaped bolalipids in each leaflet, which can result in a different U-shape fraction between leaflets and thus asymmetry between leaflets. Fig. S10 confirms U-shape head fraction asymmetry that increases with curvature, for both flexible ($k_{\text{bola}} = 0$) and moderately stiff bolalipids ($k_{\text{bola}} = 1\,k_B T$).

Together, these two effects result in membrane softening under curvature for the moderately stiff bolalipids, but constant rigidity for flexible bolalipids (Fig. 2F). In details: for membranes composed of moderately stiff bolalipid molecules ($k_{\text{bola}} = 1\,k_B T$), the U-shape bolalipid head fraction only increases in the outer leaflet, going from 10 to 20% (Fig. S10). This is in the high sensitivity region where the bending rigidity is expected to change the most (Fig. S11). We hypothesize that the molecular rigidity of a U-shaped bolalipid creates compression on the outer leaflet that stabilizes the membrane curvature and thus causes membrane softening. We suspect that for membranes composed of rigid bolalipids ($k_{\mathrm{bola}} > 1\,k_{\mathrm{B}}T$), the effect is likely not present due to the absence of U-shape formation even under strong bending.

By contrast, for membranes composed of flexible bolalipids ($k_{\mathrm{bola}} = 0$), the U-shaped bolalipid head fraction changes relatively little from its value for flat membranes (from 50% to respectively 60 and 40% for the outer and inner leaflet, Fig. S10). This is in the region where the membrane bending rigidity is expected to respond weakly to U-shape fraction (Fig. S11). Additionally, the change is symmetric, so presumably the outer leaflet becomes softer as the inner leaflet becomes stiffer, thus creating opposing effects and only weakly affecting the membrane bending rigidity as a whole. We note that the distinction between the U-shape head fraction that we plot (Fig. S10) and U-shape fraction (Fig. S11) matters little for this analysis.

### 9. DETERMINING THE GAUSSIAN BENDING RIGIDITY

In order to determine the Gaussian bending rigidity $\bar{\kappa}$ both of bilayer and flexible bolalipid membranes, we closely followed the procedure as detailed in [2]. As a consequence, we here only describe the general idea and provide the specific parameter values that we used. Details of the simulation protocol can be found in the original publication.

In short, we registered the closing frequency of membrane patches that we initialized as spherical caps. By simulating the closing of membrane patches many times and then repeating the procedure for different degrees of spherical cap opening $x$, we could sample the probability $P(x)$ for precurved caps to close. By fitting $P(x)$ with an analytical expression, the folding probability was related to the dimensionless parameter $\xi$, which itself was related to the cap radius $R$, the membrane line tension $\gamma$, the membrane rigidity $\kappa$, and the Gaussian rigidity $\bar{\kappa}$. Using all (known) parameters, we then determined $\bar{\kappa}$, following [2]

$$\bar{\kappa} = \frac{\gamma R}{\xi} - 2\kappa\,. \tag{S7}$$

For the folding simulations, we either used 1000 bilayer lipids or 500 flexible bolalipids ($k_{\mathrm{bola}} = 0$) and we simulated at $\omega = 1.5\sigma$ and $T_{\mathrm{eff}} = 1.2$ or $T_{\mathrm{eff}} = 1.3$. Initially, we distributed the lipids along a spherical cap. We allowed the lipids to reach equilibrium while we limited the available space of the lipids between two spherical shells. The full equilibration took $10250\tau$, while the last $100\tau$ were used to introduce different seeds. After the equilibration step, we run the simulation until the precurved membrane patch either fully closed or became flat, determined by the relative shape anisotropy $\kappa_{\mathrm{s}}^2$. We used the same cut-offs for $\kappa_{\mathrm{s}}^2$ as in [2]. We repeated the simulations for 8 different values of $x$ and 200 seeds each. The radius of the spherical caps $R$ was determined from the cap areas. The cap areas were determined from the averaged number of lipids per cap and the area per lipid. For the line tension, we determined the tension that acts on the open edge of a flat membrane in periodic boundary conditions, following the protocol described in [2]. Using the values for $\xi$, $R$, $\gamma$ and $\kappa$, we then determined $\bar{\kappa}$. All values are summarized in Table S4.

### 10. SIMULATION PROTOCOL FOR CARGO BUDDING

To model a spherical cargo of radius $R_{\mathrm{c}} = 8\sigma$ being adsorbed by a membrane, we set up a Lennard-Jones potential between the cargo and lipid beads using Eq. (S2). We parametrize the strength of

| Memb | $T_{\text{eff}}$ | A/lipid $[\sigma^2]$ | $R$ $[\sigma]$ | $\xi$ | $\gamma$ $[k_\text{B}T/\sigma]$ | $\kappa$ $[k_\text{B}T]$ | $\bar{\kappa}$ $[k_\text{B}T]$ | $-\bar{\kappa}/\kappa$ |
|---|---|---|---|---|---|---|---|---|
| Bila | 1.2 | 1.3587 | $7.238 \pm 0.046$ | $1.679 \pm 0.010$ | $1.237 \pm 0.035$ | $4.82 \pm 0.08$ | $-4.30 \pm 0.22$ | $0.89 \pm 0.04$ |
| Bola | 1.2 | 1.2902 | $7.165 \pm 0.000$ | $1.309 \pm 0.009$ | $1.957 \pm 0.043$ | $7.88 \pm 0.14$ | $-5.04 \pm 0.37$ | $0.64 \pm 0.04$ |
| Bila | 1.3 | 1.4238 | $7.199 \pm 0.122$ | $1.660 \pm 0.008$ | $0.917 \pm 0.031$ | $3.36 \pm 0.05$ | $-2.74 \pm 0.18$ | $0.81 \pm 0.05$ |
| Bola | 1.3 | 1.3475 | $7.322 \pm 0.000$ | $1.366 \pm 0.009$ | $1.381 \pm 0.038$ | $4.79 \pm 0.14$ | $-2.17 \pm 0.35$ | $0.45 \pm 0.07$ |

Table S4. The membrane type, $T_{\text{eff}}$, the area per lipid, the patch radius $R$, the folding parameter $\xi$, the membrane line tension $\gamma$, the bending rigidity $\kappa$, the Gaussian bending rigidity $\bar{\kappa}$, and the ratio of $-\bar{\kappa}/\kappa$.

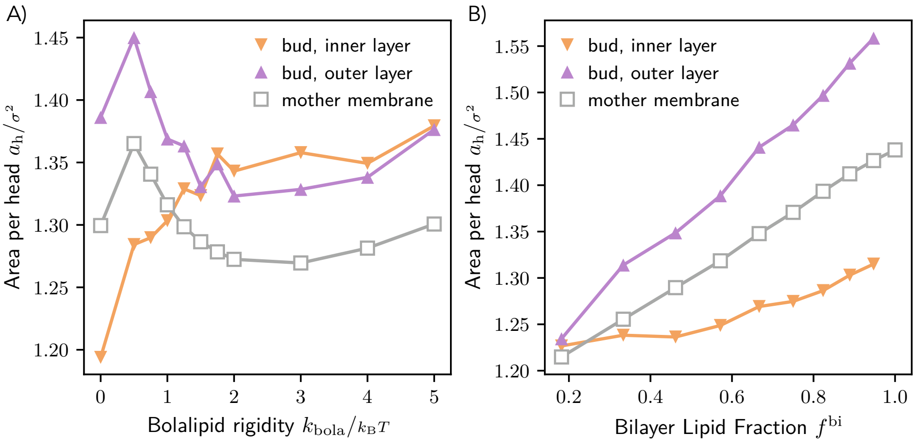


Figure S12. Area per head bead measurements, in the inner and outer layer of the membrane bud, as well as the flat mother membrane, for (A) bolalipid membranes as a function of $k_\text{bola}$ and (B) membranes of a mixture of bilayer lipids and bolalipids as a function of the bilayer lipid fraction $f^\text{bi}$.

the potential $U_m = \epsilon_\text{mc}$, calling $\epsilon_\text{mc}$ the adsorption energy. With lipid tail beads, the interaction is purely repulsive, with $r_m = r_c = 2^{1/6}(8 + 0.5)\sigma \approx 9.5\sigma$. With lipid head beads, we set up an attractive well by setting $r_c = 2^{1/6}(8 + 0.5) \cdot 1.2\sigma \approx 11.5\sigma$, giving the well a width of $\approx 2\sigma$. This limits this attractive interaction to the head beads of the membrane leaflet closest to the cargo.

For budding simulations (Fig. 4 and Fig. 5), we first pre-equilibrated for $10^4\tau$ a membrane with $60^2$ head beads per leaflet before placing the cargo bead on top of the membrane. To displace any lipids that might be inside the cargo's volume, we performed a short run in the constant volume ensemble, while scaling from zero to full strength the membrane-cargo interaction. We then ran the simulation at constant pressure until the system was in equilibrium for at least $\Delta_\text{eq} = 30 \times 10^3\tau$; the end state should then be either partial or full membrane budding. In practice total runtime was $\Delta_\text{total} \in [60, 120] \times 10^3\tau$. For a significant region of membrane parameters of interest, as we increased $\epsilon_\text{mc}$ we observed the equilibrium state transition directly from non-budded to a collapsed state with the membrane folded around the cargo in a small simulation box. To impose the existence of a region with budding we kept all membranes in budding simulations minimally stretched by setting the barostat target lateral pressure to $P_\text{xx} = P_\text{yy} = -0.001 k_\text{B}T/\sigma^3$.

Note that it was not possible to measure $\epsilon^*_\text{mc}$ for $f^\text{bi} = 1$ since budding was always followed by membrane disassembly as the bilayer membrane is close to the gas phase for the used parameters.

### 11. AREA PER HEAD MEASUREMENTS

Fig. S12A shows the area per head bead as a function of $k_{\mathrm{bola}}$ in the inner and outer layer of the membrane bud and the flat mother membrane.

We can use the area per head as a proxy for tension, by comparing with the values for the flat mother membrane. We note that these measurements excluded membrane within $3\sigma$ of a pore. The plot shows that for bolalipid membranes while the area per head is non-monotonic in $k_{\mathrm{bola}}$, it varies by less than $\approx 4\%$. In contrast, for membrane mixture of bilayer and bolalipids the area per head as a function of $f^{\mathrm{bi}}$ monotonically increases (Fig. S12B), varying by $\approx 10\%$. By default, both inner and outer leaflet would be relaxed if possible, at the same value of area per head as the flat mother membrane. However, the cargo bead competes with tension and pulls head beads from the outer to the inner layer, thus increasing the area per head for the outer layer and decreasing it for the inner layer. This happens both for relatively flexible bolalipids and mixture membranes with increasing bilayer fraction, which can indeed move head beads between leaflets, by, respectively, forming and/or flip-flopping U-shaped bolalipids, and flip-flopping bilayer lipids. When this is not possible, the area per head values for inner and outer layer are roughly equal, such as for the mixture membranes with nearly no bilayer lipids at $f^{\mathrm{bi}} \approx 0.2$. This also happens for bolalipids with $k_{\mathrm{bola}} > 2\, k_{\mathrm{B}} T$, where we note that the area per head for the bud layers are higher than that for the relaxed membrane; this can be understood by considering that at these $k_{\mathrm{bola}}$ values the bud membrane has pores, whose line tension then competes with tension, stretching the membrane. Lastly, the trend inversion for bolalipids at $k_{\mathrm{bola}} = 0.5\, k_{\mathrm{B}} T$ can be understood by considering our choice of $T_{\mathrm{eff}}$ for each $k_{\mathrm{bola}}$ changes slope at the same point (see Fig. 2A).

### 12. MEMBRANE STRUCTURE AND LIPID CONFORMATION

We developed a pipeline for analysing each frame of membrane simulation trajectories, using the pipeline framework and components from OVITO [7].

**Cluster identification and surface reconstruction** We distinguished and identified clusters by fitting mean position and orientation of lipids to the expected membranes: a horizontal plane for for the flat membrane simulations and the mother membrane in budding simulations and a sphere for the budded membrane in budding simulations. We constructed the membrane surface from the set of lipid beads using the alpha-shape method with radius $1.5\sigma$ (implemented in [7]); this ensures any pore of diameter $> 1\sigma$ will be not be closed over by the resulting surface.

**Midplane construction** We then constructed the midplane of the membrane by clustering the faces of the Voronoi diagram of the membrane surface vertices that are nearly coplanar and inside the membrane surface and then meshing the resulting oriented point cloud. This procedure is general enough to work for pre-budding frames of the budding simulations. The midplane orientation is adjusted to be consistent frame to frame.

**Membrane pore identification** By computing the signed distance to the midplane, we assigned a leaflet to each membrane surface element. We then intersected the membrane surface with the midplane surface, obtaining for each pore a line marking its perimeter. For our purposes it was sufficient to project each pore perimeter into a least-squares fitted plane and compute the area and perimeter from the projected line. For leaflet area measurements we considered the two surfaces at equal distance between the membrane surface and the midplane.

For the measurements of pore diameter in Fig. 4D and Fig. 5D, we first took the ensemble average, i.e. time average with rescaling of std. mean deviation according to autocorrelation, of the total pore area.

We consider a point to be part of a pore if its projection in the midplane membrane is within $3\sigma$ of its surface. We can also assign to a point a leaflet corresponding to its signed distance to

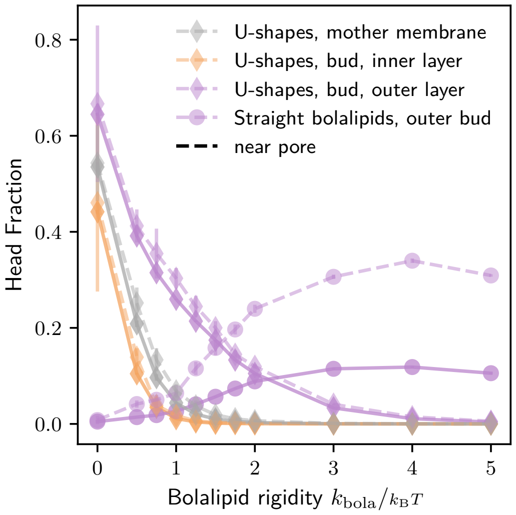


Figure S13. Lipid conformation and location in the membrane bud for pure bolalipid membranes, excluding lipids near pores (solid lines) and exclusively considering lipids near pores (dashed lines).

the midplane. We apply this location classification to the head bead of each bilayer lipid and each half of a bolalipid. Due to the bolalipid's symmetry, some combinations of conformation (given by the sign of the angle between their head beads) and location are indistinguishable, while others are transition ephemeral states that for our ensemble measures were not relevant (e.g., U-shapes that have one head bead in each layer). When both head beads are on the same leaflet, the only relevant states are the U-shape, where the heads beads angle is $\leq \pi/2$ and the flat state, where the angle is $> \pi/2$. When the head beads are in different leaflets, we get the straight state. Thus, each lipid is assigned a state, some of which have a leaflet; additionally we consider a lipid to be in a pore if any of it(s) head bead(s) is near a pore.

For simulations where pore formation was significant, namely the simulations with pure bolalipid membranes, we present in the main text conformation measurements that exclude lipids that are in a pore. To clarify that this does not qualitatively change our results, we redid the measurements in Fig. 4C, considering exclusively lipids near pores and compared (see Fig. S13). Not surprisingly, the only value to change significantly is the fraction of flat bolalipids, which is much larger near pores.

**Lipid specie/conformation per leaflet** For analysing membrane composition in lipid specie and conformation, we present head bead fractions. In this way, the denominator is a function of geometry and thus approximately conserved. For instance, the fraction of head beads that belong to bilayer lipids $f_{\mathrm{h}}^{\mathrm{bi}}$ is related in the following way to the fraction of lipids that are bilayer lipids, $f^{\mathrm{bi}}$

$$f_{\mathrm{h}}^{\mathrm{bi}} = \frac{n^{\mathrm{bi}}}{n^{\mathrm{bi}} + 2n^{\mathrm{bola}}} = \frac{1}{\frac{2}{f^{\mathrm{bi}}} - 1} = f^{\mathrm{bi}} \frac{1}{1 + (1 - f^{\mathrm{bi}})}, \tag{S8}$$

where $n^{\mathrm{bi}}$ is the number of bilayer head beads and $n^{\mathrm{bola}}$ is the number of bolalipid head beads. Therefore, near $f^{\mathrm{bi}} = 0$, $f_{\mathrm{h}}^{\mathrm{bi}}$ is nearly half $f^{\mathrm{bi}}$, while near $f^{\mathrm{bi}} = 1$, $f_{\mathrm{h}}^{\mathrm{bi}} \approx f^{\mathrm{bi}}$ .

### 13. LIPID CONFORMATION DURING BUDDING

To justify our qualitative observations of bilayer lipids and U-shaped bolalipids, respectively in mixtures and pure bolalipid membranes, enriching at the positive curvature leaflets during budding simulations, we plot the average composition of the axially symmetric profile around the cargo bud. To do this, we chose a trajectory section where the shape of the membrane was roughly static. Then within a $2000\tau$ time interval, we sampled uniformly 100 frames. For each frame, we took a cylindrical section of the simulation centred on the membrane and axis pointing upwards towards $z$. We assigned to each particle a conformation or specie, respectively. Using the cargo bead as the origin for cylindrical coordinates $(r, \theta, z)$, dropping the angle, and using hexagonal binning on the $r, z$ plane, for each conformation / species we computed the total number of particles observed. Using these totals we then thresholded on a minimum number of 40 particles observed per bin and plotted the composition as a fraction of (#studied conformation or specie) / (total in bin) (Fig. S14).

### 14. SCALING TEMPERATURE

In this section we show that little changes in our work when scaling temperature $T$ instead of our effective temperature $T_{\text{eff}}$. If we compare our phase diagrams when scaling $T_{\text{eff}}$ to scaling simulation temperature $T$, we obtain very similar results (Fig. S15).

Naturally the fraction of U-shapes becomes now a function of temperature, but by keeping to the previously defined $(T_{\text{eff}}, k_{\text{bola}})$ line we maintain the same qualitative disappearance of U-shapes as $k_{\text{bola}}$ is increased (Fig. S16). The effect on bolalipid fraction and bending rigidity becomes respectively less abrupt and strong (Fig. S17), but qualitatively we have the same trends as discussed in the main text (Figs. 2B and 2C).

### 15. MAXIMUM CURVATURE AND THE VALIDITY OF THE MONGE GAUGE

In this section we show that the maximum curvature imposed on the membrane during fluctuation spectrum measurements is not sufficient to either invalidate the use of the Monge gauge or reach the curvature-dependent regime found for semi-flexible bolalipid membranes (see Fig. 2).

We first validate the Monge gauge for describing the membrane surface. We can parametrize a periodic surface without overhangs by a height field $h(x, y)$ relative to a reference plane; we can then perform a Fourier transform in space on this field:

$$h = h(\vec{r}) = \sum_n a_n e^{i(\vec{q}_n \cdot \vec{r} + \phi_n)} = a_0 + \sum_{\substack{n \in n_+: \\ n_x > 0 \text{ or} \\ n_x = 0 \& n_y > 0}} h_n \cos(\vec{q}_n \cdot \vec{r} + \phi_n) ,$$

where $n$ are mode number vectors, and to each instantaneous real amplitude $h_n := 2a_n$ corresponds one degree of freedom. $a_0$ is just the average membrane height, kept constant by ensuring the membrane momentum is zero, via constraining the initial thermalization and the thermostat. The wave vector components are $q_{n,i} = 2\pi n_i / L_i$; we used $a_n = a_{-n}$ and for the phase $\phi_n = -\phi_{-n}$.

To apply the Monge gauge means taking several approximations on derivatives of $h$, and thus is only valid if $|\nabla h| \ll 1$. We expand to obtain:

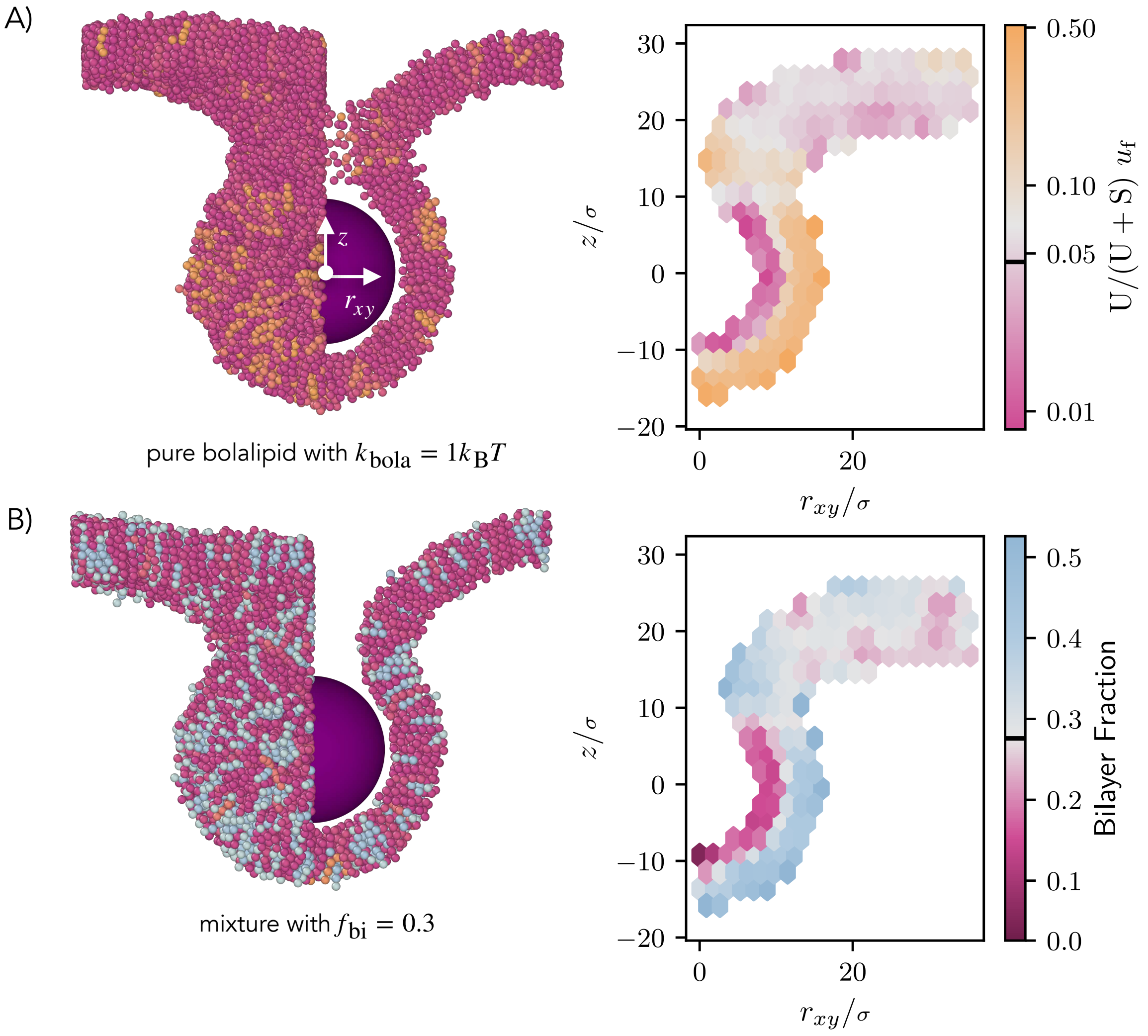


Figure S14. Lipid conformation and location in the membrane bud for pure bolalipid membranes, before budding, (A) for pure bolalipids at $k_{\rm bola} = 1\,k_{\rm B}\,T$ and for (B) a mixture of bolalipids with 30% bilayer, averaged over 100 frames spaced over a $2000\tau$ time interval. For both cases, a snapshot (left) is shown with the front right half of the membrane cut away, showing the profile shape, matching the (right) time averaged spatially varying conformation or specie fraction. Visible at $z > 10\,\sigma$ and for small $r_{\rm xy}$ is the inner region of the neck. While for mixture membranes the effect on bilayer fraction is visible using a linear scale, for bolalipid membranes at $k_{\rm bola} = 1\,k_{\rm B}\,T$ the effect spans multiple orders of magnitude so a logarithmic scale was used. The flat membrane values, taken as average values of the bins at $r_{\rm xy} > 20\,\sigma$, are indicated by a black mark on the colour bar.

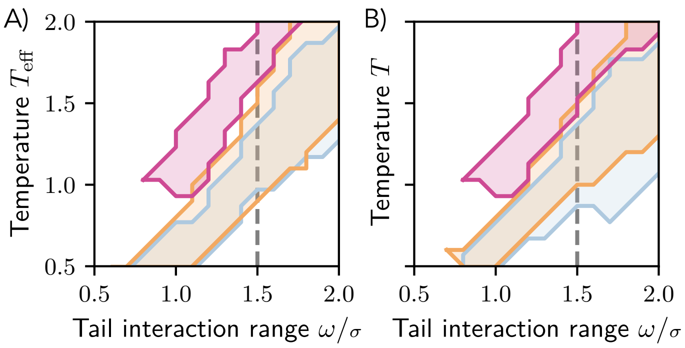


Figure S15. Phase diagram for pure bilayer and bolalipid membranes (flexible and rigid), as in Fig. 1F, for (A) $T_{\text{eff}}$ and (B) $T$ scaling.

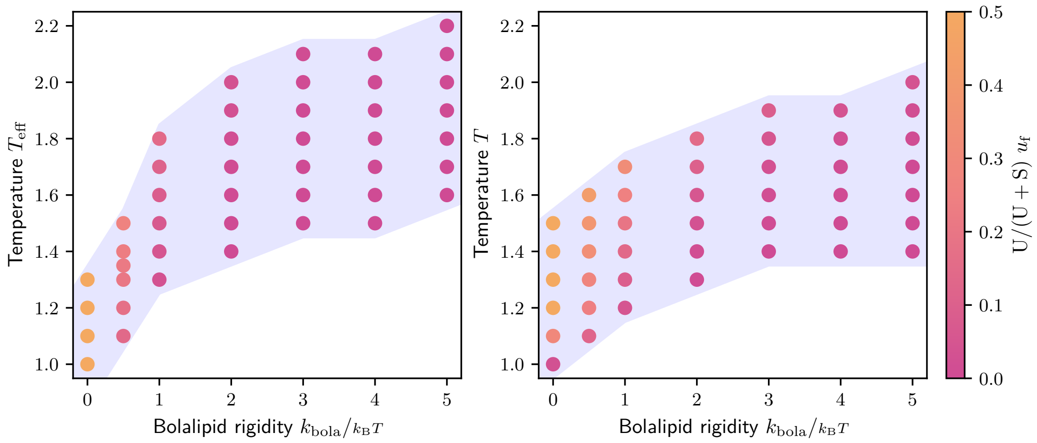


Figure S16. Liquid region (blue) for bolalipid membranes with potentials setup as in the main text as a function of $k_{\text{bola}}$ and $T_{\text{eff}}$ (A), and as a function of $k_{\text{bola}}$ and $T$ (B), with U-shape fraction in colour.

$$(\nabla h)_i = \sum_{i,n \in n_+} h_n q_{n,i} \sin\left(\vec{q}_n \cdot \vec{r} + \phi_n\right) .$$

Since we observed amplitude decreasing with $|n|$, the first two modes, with $|n| = 1$, contribute the most to the gradient. We discard the other modes and obtain an upper bound:

$$|\nabla h| \leq \frac{2\pi}{L} \sqrt{h_{1,0}^2 + h_{0,1}^2} . \tag{S9}$$

How small should be $|\nabla h|$? From Cooke and Deserno Fig. 5 (bilayer membrane with $w = 1.4$, $\epsilon = 1$), we find $L = 50$, $\langle h^2 \rangle L^2 \approx 500$ for $n = 1$ (the point x-axis coordinate matches up with expected $(2\pi/50)^2 = 0.016$). Assuming the average was taken over both modes (with $\vec{n} = (0, 1)$ and

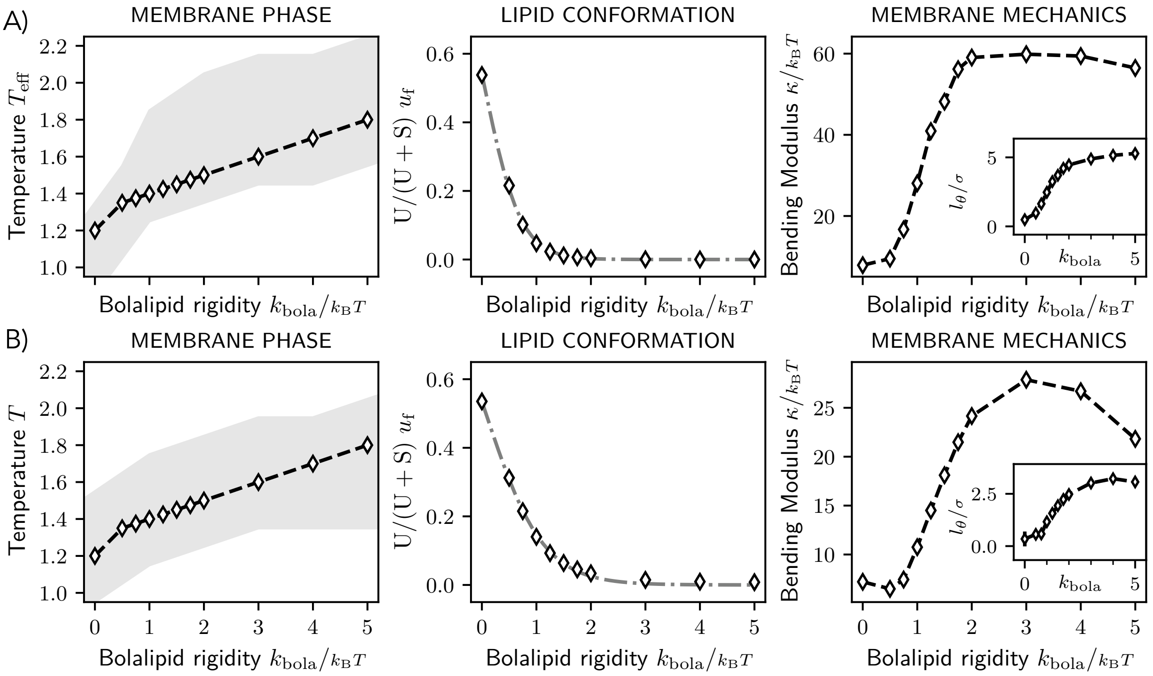


Figure S17. Comparison between using $T_{\mathrm{eff}}$ for Fig. 2 (A), with scaling temperature $T$ for the same parameters (B). The tilt contribution for $k_{\mathrm{bola}} < 1\, k_{\mathrm{B}}\, T$ is negligible.

$\vec{n} = (1, 0)$) we can write:

$$\langle h_n^2 \rangle_{|n|=1}^{1/2} = \left\langle \sqrt{\frac{h_{1,0}^2 + h_{0,1}^2}{2}} \right\rangle \leq \max \sqrt{h_{1,0}^2 + h_{0,1}^2}\ .$$

While we do not have access to the data to determine the RHS, since the LHS is strictly less or equal to the RHS we obtain that for Cooke and Deserno the threshold for the norm of the gradient was at 8% or more.

In our sims, we left the modes with $n \leq 2$ unequilibrated. Therefore, we cannot use directly averages for these modes, since they might smooth out an otherwise excessive deviation from the Monge Gauge. Instead, we compute $\max |\nabla h|$ for each frame, take the average of each $1000\tau$ block to exclude transient behaviour unlikely to affect membrane properties, and take the maximum of these averages over the full trajectory. We take the maximum over replicas.

In our simulations, we thus obtain, for the bilayer at $T_{\mathrm{eff}} = 1.1$, 12%; for flexible bolalipids ($k_{\mathrm{bola}} = 0$), 13%; for stiffer bolalipids ($k_{\mathrm{bola}} = 1\, k_{\mathrm{B}}\, T$), 6%, and for rigid bolalipids ($k_{\mathrm{bola}} = 5\, k_{\mathrm{B}}\, T$), 4%.

All of these values are close by 5% at most to the threshold used by Cooke and Deserno, so we are confident to apply the Monge Gauge. Arguably by forcing the most flexible of these membranes to stay flat (e.g. by fixing the simulation box size to a slightly larger value), we could reduce the deviation at the cost of introducing excessive tension, which would mask the effect of the bending modulus $\kappa$.

Improvements notwithstanding, continuing with the Monge Gauge, we have for the maximum mean curvature the expression:

$$\begin{aligned} H_{\max} &= \max \frac{|\nabla^2 h|}{2} \\ &= \max \frac{1}{2} \left| \sum_{n \in n_+} h_n |\vec{k}_n|^2 \cos\left(\vec{k}_n \cdot \vec{r} + \phi_n\right) \right| \\ &\leq \frac{|h_{1,0} + h_{0,1}|}{2} \left(\frac{2\pi}{L}\right)^2 , \end{aligned}$$

where in the last equality we restricted ourselves to the two largest wavelength modes.

Since we are interested in the steady state, we repeat the procedure used for obtaining the slope, obtaining, respectively, for each membrane, 0.025 (bilayer), 0.028 (flexible bolalipids), 0.012 (stiffer bolalipids) and 0.01 (rigid bolalipids), in $\sigma^{-1}$.

The maximum mean curvature for the stiffer bolalipids ($k_{\text{bola}} = 1\, k_{\text{B}}\, T$) reaches at most $0.012\sigma^{-1}$ in a fluctuation measurement. We can interpolate the data from Fig. 2F to obtain that at this mean curvature, the bending modulus is reduced by 7%, compared to its value for a flat membrane. Given this is for the maximum value of membrane mean curvature, we expect a much smaller change on the effective bending rigidity. Therefore, we find our analysis of their spectrums, which assumes a constant bending modulus, valid.

### 16. TILT MODEL VERSUS TENSION MODEL

**Membrane tension cannot explain the rigid bolalipid spectrum.** The tilt model has different asymptotic behaviour compared to the tension model. The tilt model exhibits the functional form $1/\left(\kappa q^4\right) + 1/\left(\kappa_\theta q^2\right)$. In contrast, the tension model exhibits the functional form $1/\left(\kappa q^4 + \Sigma q^2\right)$. For the tilt model, while for small $q$ the amplitude is proportional to $q^{-4}$, for large $q$ the amplitude is proportional to $q^{-2}$. In contrast, for the tension model (with positive tension) while for small $q$ the amplitude is proportional to $q^{-2}$, for large $q$ the amplitude is proportional to $q^{-4}$. If membrane tension were to be negative in the tension model, the slope would cross from negative infinity for small $q$ to $-4$ for large $q$. Consequently, between the tilt and tension models, the only model that can fit the measured slope that goes from $-4$ to $-2$ with increasing $q$ is the tilt model (best fits for each shown in Fig. S18).

Moreover, we indirectly determined membrane tension by fitting both tension and tilt with the expression:

$$\langle |h_q|^2 \rangle = \frac{k_{\text{B}} T}{L^2} \left( \frac{1}{\kappa q^4 + \Sigma q^2} + \frac{1}{\kappa_\theta q^2} \right) . \tag{S10}$$

The fit to this model with both tension and tilt overlaps with the tilt only model (also shown in Fig. S18), and the membrane tension fitted is $-0.02 \pm 0.05\, k_{\text{B}}\, T/\sigma^2$. At the longest wavelength ($q \approx 0.1\sigma^{-1}$), the tension required to match the bending term is reached when $\kappa q^4 + \Sigma q^2 \approx 0 \leftrightarrow \Sigma = -\kappa q^2 \approx -0.6\, k_{\text{B}}\, T/\sigma^2 \ll -0.02$. Therefore our barostating was effective at making tension small enough for its effect on the spectrum to be negligible.

We also measured the projected membrane tension $-L_z \langle P_x + P_y \rangle /2$, obtaining $(-2 \pm 4) \times 10^{-3}\, k_{\text{B}}\, T/\sigma^2$. This small negligible negative value of projected tension, that approximately agrees with the fitted tension, confirms our barostat was properly working.

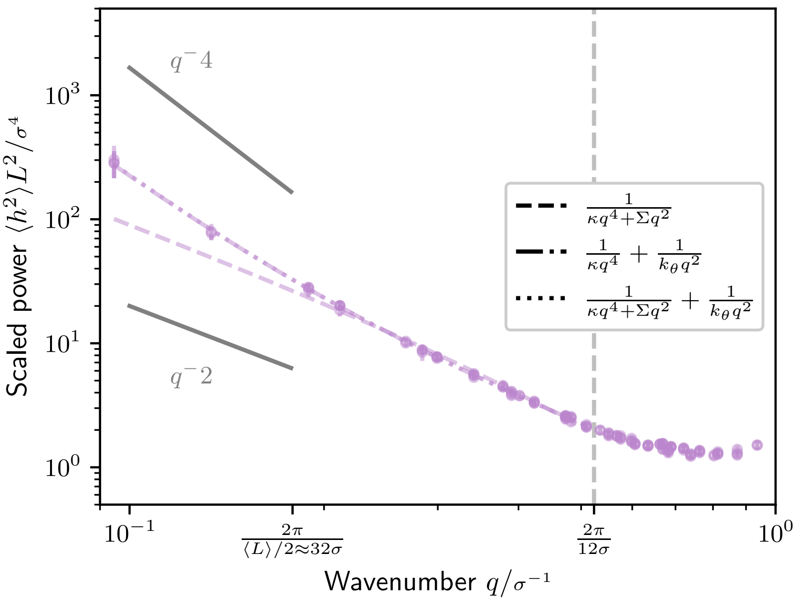


Figure S18. Height fluctuation spectrum, for a rigid bolalipid membrane at $T_{\rm eff} = 1.8$. Solid lines mark the $-2, -4$ slopes. The vertical dashed line marks the limit of the data used for fitting. The fits are different dashed lines; the fit that models both tension and tilt overlaps with the fit that only includes tilt.

### 17. EQUILIBRATION AND ENSEMBLES

Here we cover two details regarding our fluctuation simulations and analysis.

**Equilibrating the longer wavelengths does not change the relevant spectrum shown in the main text.** This is expected since our method of measurement involves averaging each mode's amplitudes over 4 replicas, and we checked that our replicas in fact were not stuck in an initial starting position and were instead exploring different regions of the phase space.

To show without doubt that this procedure does not randomly bias our results, we also ran simulations for three representative membranes until all modes were equilibrated. In order to equilibrate the long wavelength modes (mode number $|\vec{n}| < 2$), we simulated for up to $1.44 \times 10^6\,\tau$. On the all modes except the long wavelength modes ($|\vec{n}| \geq 2$), the resulting amplitudes change little (Fig. S19, 'nph+langevin (short)' and 'nph+langevin'). On the largest wavelength modes, we noticed a small deviation from theory, specifically for the bilayer and flexible bolalipid membranes ($k_{\rm bola} = 0$). These small deviations can be explained by including a negligible negative tension. Importantly, however, the resulting bending modulus $\kappa$ stays nearly the same. We note that the small negative tension disappears when we halve the timestep (Fig. S19, 'nph+langevin (timestep halved)').

**Moreover, the relevant spectrum does not depend on how the integration is done.** We tested bilayer at $T_{\rm eff} = 1.1$, flexible bolalipids ($T_{\rm eff} = 1.2, k_{\rm bola} = 0$) and rigid bolalipids ($T_{\rm eff} = 1.8, k_{\rm bola} = 5\,k_{\rm B}T$) using different integrators: zero lateral pressure with overdampened Langevin dynamics, implemented in LAMMPS via **fix nph** and **fix langevin**, as done in most of our work; a NPT ensemble, implemented in LAMMPS via **fix npt**; and Brownian dynamics in a fixed volume, implemented in LAMMPS via **fix nve** and **fix langevin**, where the box size is set to the average value in **nph+langevin** simulations. The resulting measurements of $\kappa$ are at most within 10% of

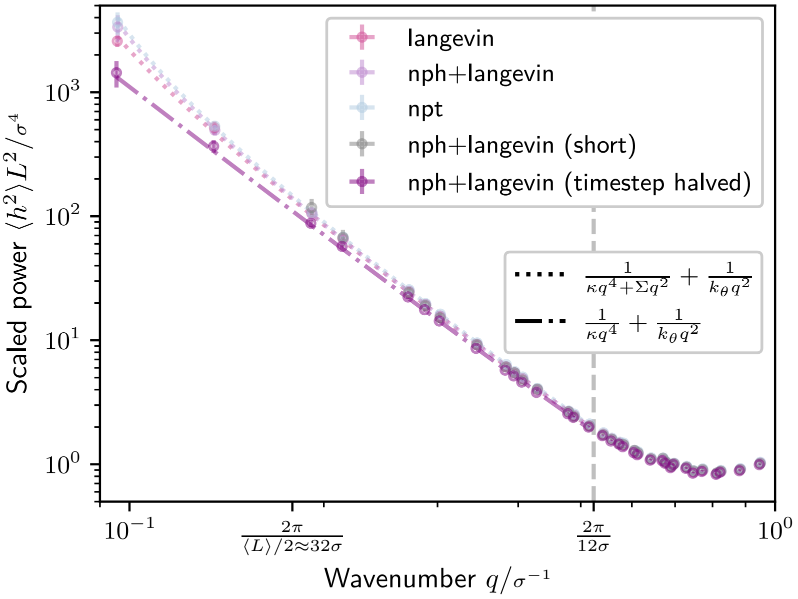


Figure S19. Height fluctuation spectrum, for a bilayer membrane at $T_{\text{eff}} = 1.1$ simulated under different integrators. The resulting fit parameters together with the normalized $\chi^2$ are shown in Tabl. S5.

the values reported in the main text (Tabl. S5). The value of $l_\theta$ does increase by 30% for this bilayer system, but since it is still below our main text threshold of $2\sigma$ our conclusions do not change.

To start with, we get the same results if we equilibrate first with a barostat, fix the box size, and only then sample the fluctuation spectrum (Fig. S19,'langevin'). It becomes then easy to expect that there is no significant difference for our simulations between using **fix nph** and **fix langevin** (Fig. S19,'nph+langevin') versus **fix npt** (Fig. S19,'npt'). Our barostat simply equilibrates our membrane to near zero tension and then does not significantly contribute to the system. While for bilayer and flexible bolalipid membranes, the first modes (i.e. $|\vec{n}| < 2$) amplitudes are noticeably above the fitted line for the tilt-including model, these can be reached by including a negative tension term with the negligible value of $-0.05\, k_{\text{B}}T/\sigma^2$ (Fig. S19,'npt' and 'nph+langevin'). We call this value negligible, since it is only roughly 3% of the tension necessary to rupture a membrane in this model [1], and its effect is long vanished at the wavelengths we use for the fits in the main text. We understand this not as an issue with our barostating, but simply as consequence of the $q^{-2}$ dependency on tension. Sufficiently large membranes and small bending modulus $\kappa$ will eventually reveal tension present either due to integration error or the approximation of membrane tension by the projected membrane tension. Nevertheless, to show nothing changes in the spectrum used in the main text, we halved the timestep (setting $\Delta t = 0.005\,\tau$), which reduces the fitted tension by 3x and allows a good fit over the entire spectrum with the model that only includes tilt and not tension (Fig. S19,'timestep halved'). For rigid membranes (e.g. rigid bolalipid membranes), all modes are equilibrated for the shorter duration of $60 \times 10^3\,\tau$ used in the main text results, and we obtain the same results regardless of the integration scheme used (npt, nph+langevin, or fixed box size with Langevin).

| | $\chi^2/N$ | $\kappa/k_B T$ | $\Sigma/\left(k_B T/\sigma^2\right)$ | $l_\theta/\sigma$ |
|---|---|---|---|---|
| langevin | 0.72 | 8.51±0.12 | -0.039±0.004 | 0.85±0.05 |
| nph+langevin | 0.28 | 8.71±0.11 | -0.050±0.004 | 0.92±0.04 |
| npt | 0.25 | 8.35±0.09 | -0.0492±0.0026 | 0.86±0.04 |
| nph+langevin (short) | 0.17 | 7.88±0.25 | N/A | 0.60±0.16 |
| nph+langevin (timestep halved) | 0.31 | 9.13±0.12 | N/A | 0.97±0.05 |

Table S5. Normalized $\chi^2$ and parameters for fits in Fig. S19.

### 18. SUPPLEMENTAL MOVIES.

**Movie S1** Self-assembly of flexible bolalipid molecules into a flat membrane ($k_{\text{bola}} = 0$).

**Movies S2** Close-up view of a flat membrane, with tail beads shown like sticks.

- **Movie S2.a** Membrane made of bilayer-forming molecules at $T_{\text{eff}} = 1.1$ (same bilayer membrane as in tethers of Fig. 2F)
- **Movie S2.b** Membrane made of flexible bolalipid molecules ($k_{\text{bola}} = 0$).
- **Movie S2.c** Membrane made of rigid bolalipid molecules ($k_{\text{bola}} = 5\, k_B T$).

**Movie S3** Fluctuating flat membrane patch of bolalipid molecules at $k_{\text{bola}} = 0$ used for height fluctuation spectrum measurements.

**Movie S4** Cylindrical membranes after equilibration, made of bolalipid molecules of intermediate stiffness ($k_{\text{bola}} = 1\, k_B T$) at the largest simulated radius $R \approx 17\sigma$.

**Movies S5** Successful budding, showing only a cargo-centred cross-section.

- **Movie S5.a** Budding from a membrane made of flexible bolalipid molecules ($k_{\text{bola}} = 0$).
- **Movie S5.b** Budding from a membrane made of bolalipid molecules of intermediate stiffness ($k_{\text{bola}} = 1\, k_B T$).
- **Movie S5.c** Budding from a membrane made of completely stiff bolalipid molecules ($k_{\text{bola}} = 5\, k_B T$).
- **Movie S5.d** Budding from a membrane made of a mixture of bolalipid and bilayer-forming lipid molecules, at bilayer fraction $f^{\text{bi}} = 0.46$.

**Movie S6** Rotating snapshot of a bud formed from a membrane made out of stiff rigid bolalipid molecules ($k_{\text{bola}} = 5\, k_B T$), showing several pores ($\epsilon_{\text{mc}} = 3.25\, k_B T$).

---
